# Widespread decline in plant diversity across six decades

**DOI:** 10.1101/2020.08.31.275461

**Authors:** Eichenberg David, Diana E. Bowler, Bonn Aletta, Bruelheide Helge, Grescho Volker, Harter David, Jandt Ute, May Rudolf, Winter Marten, Jansen Florian

**Affiliations:** German Centre for Integrative Biodiversity Research (iDiv), Halle-Jena-Leipzig, Deutscher Platz 5e, Leipzig, 04103, Germany; Department of Ecosystem Services, Helmholtz Centre for Environmental Research - UFZ, Permoserstr. 15, 04318 Leipzig, Germany; Institute of Biodiversity, Friedrich Schiller University Jena, Dornburger Str. 159, 07743 Jena, Germany; Institute of Biology/Geobotany and Botanical Garden, Martin Luther University Halle-Wittenberg, Am Kirchtor 1, 06108 Halle (Saale), Germany; Federal Agency for Nature Conservation (BfN), Konstantinstr. 110, 53179 Bonn, Germany; Faculty of Agricultural and Environmental Sciences, University of Rostock, Rostock, Germany

**Keywords:** biodiversity change, floristic turnover, heterogeneous data, occurrence records, macroecology, reporting bias, spatio-temporal analysis

## Abstract

Based on plant occurrence data covering all parts of Germany, we investigated changes in the distribution of 2146 plant species between 1960 and 2017. We analyzed 29 million occurrence records over an area of ∼350.000 km^2^ on a 5 × 5 km grid using temporal and spatio-temporal models and accounting for sampling bias. Since the 1960s, more than 70% of investigated plant species showed significant declines in nation-wide occurrence. Archaeophytes (species introduced before 1492) most strongly declined but also native plant species experienced severe declines. In contrast, neophytes (species introduced after 1492) increased in their nation-wide occurrence but not homogeneously throughout the country. Our analysis suggests that the strongest declines in native species already happened in the 1960s-80s, a time frame in which usually few data exist. Increases in neophytic species were strongest in the 1990s and 2010s. Overall, the increase in neophytes did not compensate for the loss of other species, resulting in a decrease in mean grid-cell species-richness of -1.9% per decade. The decline in plant biodiversity is a widespread phenomenon occurring in different habitats and geographic regions. It is likely that this decline has major repercussions on ecosystem functioning and overall biodiversity, potentially with cascading effects across trophic levels. The approach used in this study is transferable to large-scale trend analyses using heterogeneous occurrence data.

## 1. Introduction

Biodiversity loss is one of the greatest global challenges. Approx. 1 Mio species are threatened by extinction (Diaz et al 2019). While the widespread decline of fauna (Dirzo et al. 2014) is discussed prominently in the scientific community and the general public, especially for terrestrial insects (e.g. Eisenhauer et al., 2019; Hallmann et al., 2017; Powney et al., 2015; van Klink et al., 2020), large-scale changes on the distribution of plants are less widely recognized. There are few examples of assessments of temporal trends of plant diversity over larger regions, such as whole countries (Finderup Nielsen et al., 2019; Walker & Preston, 2006). A deeper understanding of biodiversity change in plants is essential for predicting ecosystem-wide changes, including effects on ecosystem functioning (Hejda & de Bello, 2013) and the provision of ecosystem services relied on byhumans (Guo et al. 2010).

An increasing amount of studies focus on trends in plant biodiversity at global (e.g. Feeley et al., 2020; Gonzalez et al., 2016; Vellend et al., 2013; Winter et al,. 2009), regional and trans-regional scales (e.g. Bruelheide et al., 2020; Jansen et al., 2020; Finderup Nielsen et al., 2019; Jandt et al., 2011; Staude et al., 2020) and at small-scales, e.g. on the plot-level (e.g. Diekmann et al., 2014; Hüllbusch et al., 2016). However, these studies do not show consistent trends in species-richness, with findings of increases (Finderup Nielsen et al., 2019) and decreases (Meyer et al., 2013) and also of no net change (Vellend et al., 2013). Some studies indicate that trends in plant species-richness may depend on the native status of plants in the region under study (c.f. Cardinale et al., 2018). As pointed out by Sax and Gaines (2003), net gains or losses of native species in large study regions can be balanced or even overcompensated by the introduction of non-native species. For example, in a study on the flora of Denmark, Finderup Nielsen et al. (2019) reported an increased naturalization and subsequent spread in exotic species and increases in widespread natives species, but declines in rare natives over the last 140 years. In total, this led to a net gain in species-richness in the studied regions, while species communities became more homogeneous across the study regions. A study by Winter et al. (2009) showed similar results on the European scale. A Focus only on total species-richness may lead to the conclusion that biodiversity in a region remains stable, whereas it actually is experiencing turnover in species composition. This turnover may drastically alter the floristic composition in a study region – and potentially affect ecosystem functioning and stability (Naeem et al., 2012).

Analysis of floristic changes at large scales is challenged by the lack of repeated surveys of large regions (Walker & Preston, 2006, but see Switzerland and the UK, Hintermann et al., 2000, Wood et al., 2017). Typically, species-level changes at large scales, such as national or global-levels, are assessed via red lists, during structured monitoring projects. Data for the more common or non-threatened or non-iconic taxa are mostly recorded through atlas projects which do not aim at repeated recording. Consequently, the different kinds of data on plant occurrences for large regions, e.g. countries or continents come with different protocols and methods for data collection or study focus. This often leads to heterogeneous data quality and structures, hampering the analysis of such data for temporal trends. Therefore, the integration of spatially and taxonomically comprehensive data across different sources of information – often including citizen science data – has become increasingly relevant for the assessment of biodiversity change across larger spatio-temporal scales (Isaac et al., 2014, 2020; Zipkin et al., 2019).

In Germany, the majority of knowledge on changes in plant biodiversity comes from plant community resurvey studies of (semi)permanent relevés. Conclusions are mixed, showing strong (Wesche et al., 2012), little (e.g. Jensen et al., 2012) or no significant changes in local species-richness (e.g. Bruelheide, 1998; Diekmann et al., 2014; Litza & Diekmann, 2017), depending on the investigated habitats, species groups, plot sizes or the temporal extent of the study. Most studies report that declines in species-richness and changes in community composition are strongest in agricultural landscape (e.g. Meyer et al., 2013; Meyer et al., 2015). However, plot-level analyses may not allow for a straightforward extrapolation to larger scales due to several biases, e.g. in the spatial (i.e. habitat) representativeness of the plots or local differences in disturbances or management (Cardinale et al., 2018). Ignoring the local, and potentially spatially biased, small-scale patterns may lead to erroneous conclusions on large-scale net changes in biodiversity (Cardinale et al., 2018; Gonzalez et al., 2016; Sax & Gaines, 2003). Moreover, to understand changes in large-scale biodiversity it is crucial to evaluate as many species as possible.

Germany, organized in 16 federal states, offers an ideal example to demonstrate how heterogeneous datasets on plant biodiversity can be combined and analyzed to detect changes in large-scale species distributions and, subsequently, species-richness. Each federal state has carried out at least one floristic atlas survey (Bergmeier 1992) followed by other survey projects, but protocols, time-frames and predefined species lists at least for the latter varied in and across states. Although the atlas surveys have not been systematically repeated, several local and often taxonomically less comprehensive, surveys exist across Germany. The combination of this data has led to publication of the German Atlas of Vascular Plants (Netzwerk Phytodiversität Deutschland & Bundesamt für Naturschutz, 2013). However, as the underlying data often show spatial and temporal inconsistencies in the sampling coverage of subregions within the complete study region, detection of changes in biodiversity over time can be very challenging (Hill, 2012; Pescott et al., 2019).

Relevé data on the plot level are available from research institutes, universities, online databases (e.g. GVRD, Jandt & Bruelheide, 2012 or vegetweb, Jansen et al., 2015) and private collections. These data mostly have accurate geographic information, are restricted in size (one to a few square meters) and often lack temporal replication and are thus often not suitable for large-scale temporal trend analyses on their own (Chytrý, et al., 2014). Compiling and integrating different datasets (atlas, relevé and observational data from private observations, excursions, museum records, mobile apps and from spatially referenced legacy collections) in a common analysis – and thus making use of the merits of each of these data-types – may allow to quantify long-term changes of plant species distributions on large spatial and temporal scales, potentially also at a fine spatial grain. Meanwhile, modern statistical tools allow to incorporate different datatypes from different sources in robust analyses, while accounting for their heterogeneity (Isaac et al., 2014).

In the present study, we compiled an extensive dataset of the spatio-temporal occurrence of vascular plants in Germany. The dataset was collated from multiple sources on plant occurrence records and vegetation surveys in Germany, varying in taxonomic extent and sampling effort. After accounting for incomplete species recording across space and time, we a) assess spatio-temporal changes in the occurrence of 2136 species on a 5 × 5 km grid-cell basis, and b) assess the balance between winners and losers. We c) explore the temporal dynamics of these changes based on floristic status (natives vs. non-natives). Using spatio-temporal models, we d) assess changes in mean grid-cell species-richness across the whole nation, accounting for spatio-temporal dependence in the data. Moreover, we e) explore the spatial heterogeneity in the patterns of changes in grid-cell species-richness over the last six decades and identify hotspots of biodiversity turnover.

## 2. Material and Methods

We developed a workflow to harmonize the available data into a common format, taxonomically and spatially (Electronic appendix, Figure S1). We accounted for the potential effects of imperfect detection, using the Frescalo algorithm (Hill, 2012). This algorithm has specifically been developed for repeated, large-scale surveys, such as atlases, and computes species occurrence probabilities across periods of data availability (possibly irregularly spaced, Pescott et al., 2019) on a defined grid-size. However, in its original form, this algorithm does not allow for spatio-temporally explicit analyses. To this end, we further demonstrate the use of new methods to account for temporal and spatio-temporal autocorrelation in the resulting data, thus enabling robust analyses of changes in species occurrences and richness across space and time.

### 2.1. Data compilation and taxonomic harmonization

We compiled an extensive dataset of approx. 29 million occurrence records in Germany between 1960 and 2017 from 23 different data sources (Electronic appendix, Table S1). The full dataset comprises the unaggregated data underlying the German Distribution Atlas of Ferns and Flowering Plants (Netzwerk Phytodiversität Deutschland & Bundesamt für Naturschutz, 2013), restricted to observations between 1960 and 2013. This dataset – called the FlorKart dataset (Bundesamt für Naturschutz, https://www.bfn.de/themen/artenschutz/erfassung-und-kartierung/florenkartierung.html) – is a compilation of occurrence records gathered from, *inter alia*, the Floristic Atlases of Western and Eastern Germany (Benkert et al., 1996; Haeupler & Schönfelder, 1989) and floristic data extracted from 74 mapping projects. We extended this dataset to the year 2017 by integrating data from more recent habitat mapping projects of federal states, vegetation relevés provided in two major German databases, GVRD (Jandt & Bruelheide, 2012) and vegetweb 2.0 (Jansen et al., 2015) and from universities and private collections. For all datasets, observations were georeferenced on a grid-cell level (a quadrant of German ordinance maps, “MTBQ”, approx. 5 × 5 km). In all spatially explicit analyses, central coordinates of each grid-cell (in UTM) were used as sampling locations (Zuur & Ieno, 2017).

Taxa were harmonized using a common taxonomic reference list (GermanSL, Jansen & Dengler, 2008, version 1.4, https://germansl.infinitenature.org/). Subspecies, variants etc. were raised to the species or, if necessary, to the aggregate level. For simplicity, we will refer to these as “species” in the following. Taxonomic harmonization was achieved using the R-package vegdata (Jansen & Dengler, 2010). This resulted in data on the occurrence of 2976 vascular plant taxa, equaling approx. 77% of all German vascular plant taxa (5256 taxa, including subspecies and variants or 3868 taxa when raised to the taxonomic level used in this study). For analyzing trends, the dataset was binned into three periods (1960-1987, 1988-1996, 1997-2017), each of them with similar number of total records and covering all 12024 German grid-cells. The temporal extent of these periods was determined by the need to find periods of similar coverage (spatial and taxonomically) of the whole nation (see Electronic appendix, Technical details). We excluded species that were recorded in only one of the three periods or had fewer than 23 records in total (i.e. the 10% quantile of species frequency distribution data). Species were characterized according to their floristic status either as natives, archaeophytes (non-natives introduced before 1492) or neophytes (non-natives introduced after 1492), using information available from the database BiolFlor (Klotz et al., 2002) and FloraWeb (Bundesamt für Naturschutz, http://www.floraweb.de/). Species with an unknown floristic status were excluded (see all species-level details in Electronic Appendix, Table S2). This left us with a total of 2234 species for analysis, equaling 58% of all vascular plant species known from Germany (harmonized to the taxonomic level of the present analysis).

### 2.2. Correction for false absences

The majority of the data originated from grid-based occurrence-only records (approx. 95%, Electronic appendix, Table S1). In cases where occurrence records do not originate from a project focusing on the complete floristic inventory or does not use complete check-lists, false absences (i.e. not reporting a species that was present, but was either not detected or detected but not reported) are an issue (Pescott et al., 2018). In addition, atlas projects, which aim at taxonomic completeness may not be finished in a federal state completely within one of the defined study periods, leading to taxonomic or spatial gaps in the data of a single study period. To correct for this so-called reporting bias, we used the Frescalo algorithm (Hill, 2012) available in the R package ‘sparta’ (August, 2015, v. 0.1.48). Briefly, the Frescalo algorithm calculates the occurrence probability (OP) of a species not detected or reported in a focal grid-cell, based on the frequency of this species in the local neighborhood (here: 100 grid-cells) of this cell, while accounting for the ecological similarity of the neighborhood (Hill, 2012). Ecological similarity of the neighboring grid-cells was calculated based on a set of 76 variables, comprising climatic, topographic and edaphic measures. A detailed description of the specifications for the Frescalo algorithm is given in Electronic appendix (Technical details). Correction for false absences increased the dataset to more than 42 Mio entries of occurrence probabilities per species and grid-cell between 1960 and 2017.

### 2.3. Calculation of species-specific occurrence across space and time

Maps of the spatial distribution of a given species across the study region at a given time are not a direct outcome of the Frescalo algorithm as available in ‘sparta’. However, the spatial distribution of the probability of a species being present at the focal grid-cell of neighborhood in a certain period can be readily calculated from the available output using Equation 1 (c.f. Bijlsma, 2013). 

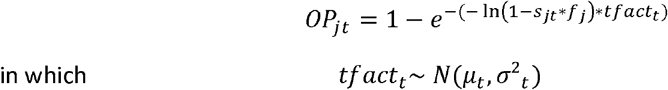

Equation 1: Calculation of species occurrence probability OP_jt_ of a species in the focal grid-cell j of a neighborhood at time t_;_ s_jt_ = sampling intensity (a measure of sampling completeness calculated by the Frescalo algorithm) for grid-cell j at time step t, f_j_ = the observed frequency of the respective species in neighborhood j, and tfact_t_ = the time factor (the estimated relative frequency of the respective species; a parameter derived by the Frescalo algorithm) at time t. OP_jt_ was calculated for each species, separately. All variables are given in the Frescalo output file provided in R (for the respective R-Script see Appendix I).

To account for uncertainty in the Frescalo estimates, calculations of species-specific occurrence probability OP_jt_ were based on 1000 realizations of μ_t_ (sampled from a species-specific normal distribution with mean μ_t_ and s_t_, c.f. Equation 1). For each species and each realization, we calculated the nation-wide occurrence of a species as the sum of all occurrence probabilities of a species (SOP_Spec_) across Germany for each period according to Equation 2. 

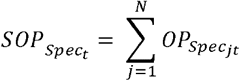

Equation 2: Calculation of nation-wide occurrence of a species (SOP_Spec_) at time t; with: Spec= species under consideration; j= index of grid-cell, t= index of timestep (i.e. 1=1960-1987; 2= 1988-1996; 3= 1997-2017); OP_Specjt_= occurrence probability of species in grid-cell j at time t (Equation 1).

### 2.4. Calculation of grid-cell species-richness

We summed up the occurrence probabilities of all species within a grid-cell as an estimate of species-richness (SOP_Grid_), while acknowledging that it is not species-richness *per-se* since our analysis does not include the very rare species (see section 2.1, 2.2 and discussion). Hence, our species-richness values are underestimating actual grid-cell species-richness. However, SOP_Grid_ was found to be significantly correlated (r= 0.39, p< 0.001, Electronic Appendix, Figure S3) to species numbers in grid-cells of the FlorKart dataset (c.f. section 2.1) that Kühn et al. (2006) identified to be well-sampled. Therefore, changes in SOP_Grid_ can be interpreted as meaningfully representing relative changes between grid-cells and timesteps. 

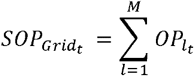

Equation 3: Calculation of grid-cell species-richness as the sum of occurrence probability (SOP_Grid_). With: l= index of the species, OP_lt_= occurrence probability of this species l in a grid-cell at time t (as derived from Equation 1).

### 2.5. Changes in species-specific occurrence

Changes in species-specific occurrences over time were evaluated at the species-level, using a Bayesian log-linear mixed effects model and including a random walk of order 1 (“rw1”) to account for temporal dependency. These specifications ensured that changes are bounded to -100% at the lower end, but not at the upper end. 

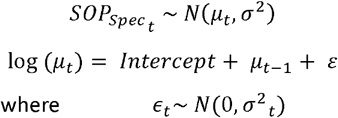

Model 1: Estimation of changes in species-specific occupancy over time according to a random walk component of order 1.

Of the total of 2234, 2206 species were found to have significant changes. A critical inspection showed that 102 species exhibited extreme trends (i.e. above or below the 95% quantile range of change). These extreme changes were discussed in depth with taxon experts, and those considered to be unrealistic trends (70 species) were omitted from further analyses (c.f. Electronic appendix, Table S3).

### 2.6. Changes in mean grid-cell species-richness

To analyze changes in nationwide mean grid-cell species-richness, we ran spatio-temporally explicit models, based on gamma distributions (selected residuals were biased in time and space for the log-normal case). To account for spatial and temporal dependencies, we included a temporal correlation structure with a random walk of order 1. Moreover, we accounted for spatial autocorrelation with an autoregressive component of order 1 (c.f. Model 2). For technical details on the configuration of the spatial component (including priors) see Electronic appendix (Technical details). 

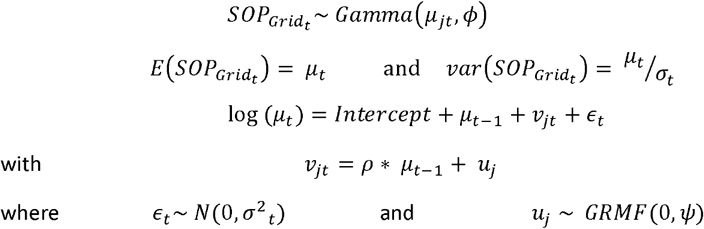

Model 2: Estimation of changes in mean grid-cell species-richness. The spatial component (v_jt_) is correlated across time with parameter *ρ* and connected to a spatial variable (u_j_), according to a Gaussian Markovian Random Field (GMRF) with mean 0 and the covariance matrix of the grid-cells (*ψ*). *ψ* is determined by the stochastic partial differential equation (SPDE) approach introduced by Lindgren, Rue, & Lindstroem (2011).

For computational reasons, analyses of spatio-temporal changes were only based on the mean OPs of the 1000 realizations per species, grid-cell and time (c.f. section 2.3.). We analyzed changes within species assemblages for each floristic status and across all species. Model predictions were used to visualize the spatio-temporal variability in changes of species-richness. Based on these values, we defined hotspots in biodiversity change as the lower and upper 10^th^ percentile of relative changes of a grid-cell between the last and first study period. Spatial correlations among these hotspots were assessed using a modified F-Test, accounting for the spatial structure in the data; post-hoc pairwise comparisons were based on Bonferroni-Holm corrected pairwise modified t-Tests. Both routines are available from the R package ‘SpatialPack’ (Vallejos et al., 2018).

For all model parameters, we used penalized complexity priors (Simpson et al., 2017) with a scaling parameter U=1 and α=0.5, ensuring an uninformative prior expectation for all model hyperparameters. All statistical analyses were carried out in R (R Development Core Team, Version 3.5.2), using the package INLA (Rue & Martino, 2009, Version 18.07.12), except the assessment of correlations between hotspots. Model residuals were visually checked for normality; for model evaluation see Electronic appendix (Technical details). In all analyses, effects were interpreted as statistically significant if the 95% credibility intervals (CI) of the (differences in) the estimated posteriors of the predictor did not overlap with zero.

## 3. Results

### 3.1. Changes in species-specific occurrences

Figure 1 shows the relative differences in species occurrences between the first (1960-1987) and the last (1997-2017) study period. Of all 2136 species, 1526 species (approx. 71%) showed a decrease and 610 species (approx. 29%) an increase in occurrence. No species showed a decline of -100%, i.e. went extinct in Germany. A detailed list of the winners and losers in occupancy for all three timesteps is given in Table S3 and in an interactive plot (https://shiny.idiv.de/de25geka/WinnersLosers/).

**Figure 1:**
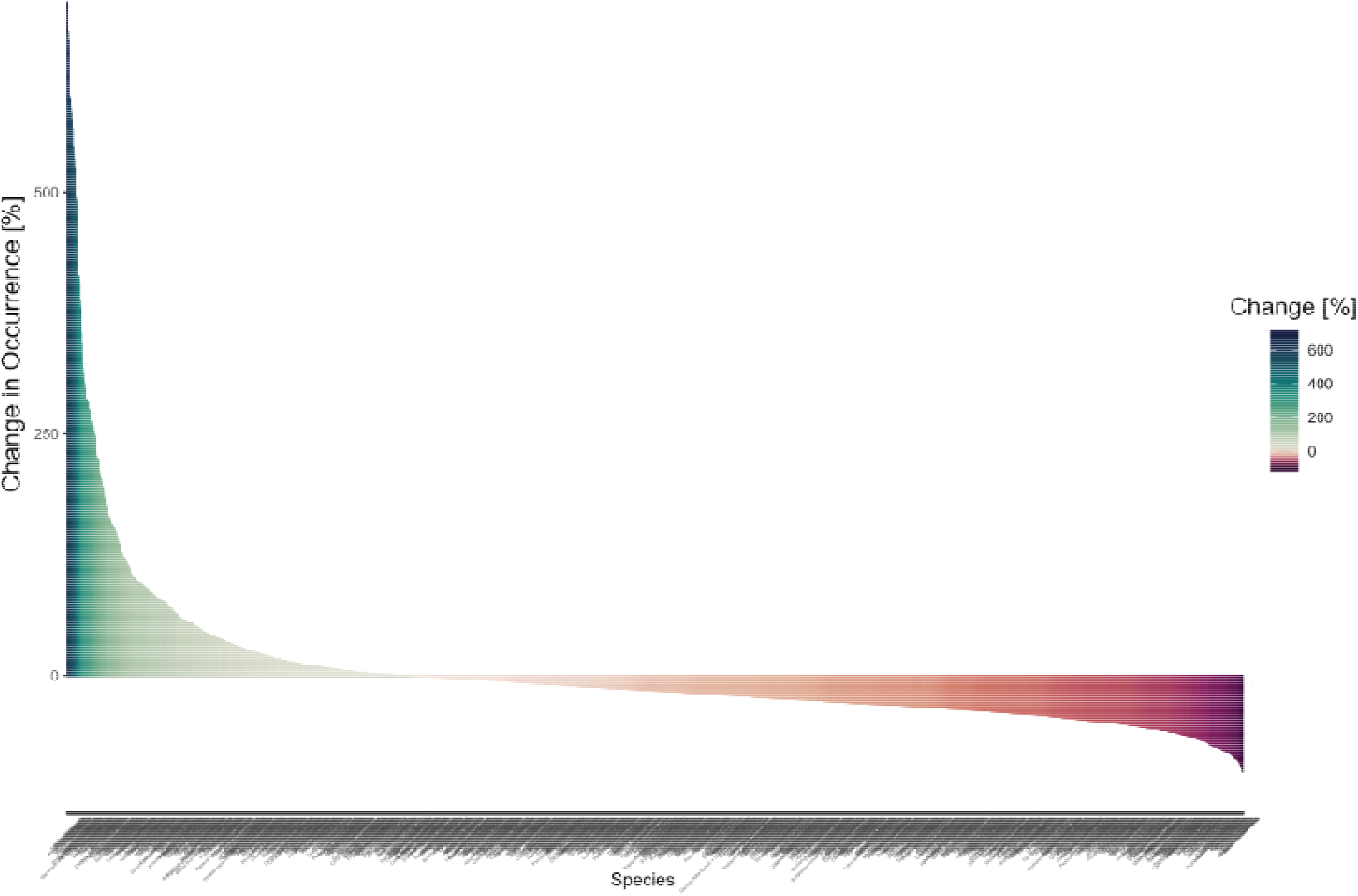
Change in occurrence in percent between 1960-1987 and 1995-2017. X-axis shows species names. Winners are shown in green, losers are depicted in red. For details in changes of species occupancy over time see Electronic appendix, Table S3. For increased visibility of species names see *https://shiny.idiv.de/de25geka/WinnersLosers/*

The species with strongest decline in mean occurrence probability (−99.8%) was *Anagallis tenella* L., a strongly endangered native species of nutrient-poor fens and transition mires in Germany. In contrast, the neophyte *Senecio inaequidens* DC., showed the strongest increase (+696%, Figure 2).

**Figure 2:**
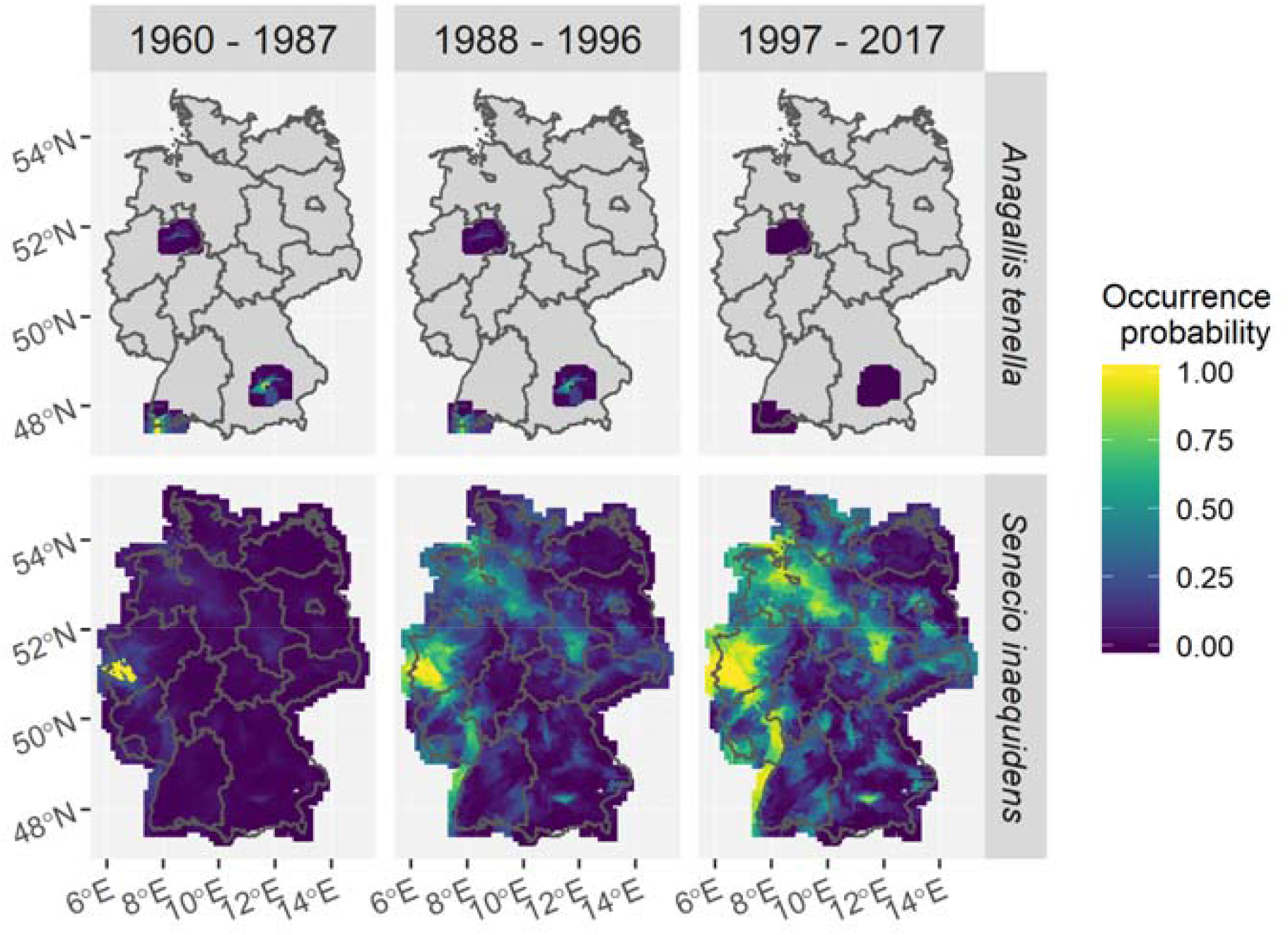
Occurrence probability estimates for the three study periods on a 5 × 5 km grid. Top: the strongly endangered native species Anagallis tenella L.. Bottom: the neophyte Senecio inaequidens DC. Gray areas are outside of the range supported by the Frescalo algorithm.

### 3.2. Changes in occurrences of floristic status groups

We found significant changes in mean occurrence among all grid cells in Germany for all floristic status groups (Table 1). Across all species, we found a total decline of approx. -10.8% over the last six decades, with strongest losses occurring between the first (1960-1987) and the second (1988-1996) study period. While natives and archaeophytes significantly decreased over the whole study period, neophytes showed a significant increase in mean occurrence. Native species and archaeophytes showed most change between the first and second periods.

**Table 1:** Changes in mean occurrence as well as relative differences [%] (incl lower and upper 95%CI) across the three study periods as predicted for the average species. Letters indicate significant differences in mean values based on the effect of the temporal component in Model 1. Asterisks indicate significant mean decadal changes. n_Specs_: number of species in each floristic status group.

| Floristic status | Absolute mean occurrence<br>(measured as $SOP_{\text{spec}}$) | | | | | | Relative Differences [%] | | | | | | Mean decadal<br>change [%] |
| --- | --- | --- | --- | --- | --- | --- | --- | --- | --- | --- | --- | --- | --- |
|  | t1 = 1960 - 1987 |  | t2 = 1988 - 1996 |  | t3 = 1997 - 2017 |  | t2 - t1 |  | t3 - t2 |  | t3 - t1 |  |  |
| Natives<br>($n_{\text{spec}} = 1724$) | 1688.7 *a* | | 1504.3 *b* | | 1459.1 *c* | | -10.9 | | -3.0 | | -13.6 | | -2.39 * |
|  | 1560.3 | 1827.8 | 1390.2 | 1628.0 | 1348.2 | 1579.2 | -10.9 | -10.9 | -3.0 | -3.0 | -13.6 | -13.6 |  |
| Archaeophytes<br>($n_{\text{pec}} = 186$) | 1301.0 *a* | | 1077.8 *b* | | 1020.1 *b* | | -17.2 | | -5.3 | | -21.6 | | -3.79 * |
|  | 1004.5 | 1687.7 | 834.4 | 1393.7 | 789.1 | 1321.2 | -17.4 | -16.9 | -5.4 | -5.2 | -21.7 | -21.5 |  |
| Neophytes<br>($n_{\text{pec}} = 226$) | 531.6 *a* | | 512.8 *a* | | 655.0 *b* | | -3.6 | | 27.9 | | 23.3 | | 4.09 * |
|  | 375.8 | 752.8 | 362.7 | 725.4 | 462.9 | 929.1 | -3.6 | -3.5 | 27.6 | 28.1 | 23.2 | 23.4 |  |
| All Species<br>($n_{\text{pec}} = 2136$) | 1510.3 *a* | | 1344.8 *b* | | 1346.6 *b* | | -11.0 | | 0.1 | | -10.8 | | -1.9 * |
|  | 1396.2 | 1633.8 | 1243.5 | 1454.5 | 1245.0 | 1456.5 | -11.0 | -10.9 | 0.1 | 0.1 | -10.9 | -10.8 |  |

Color codes: Absolute occurrence from green (high) to yellow (low); Relative differences from blue (increase) over grey to brown (decrease).

Among the floristic status groups, archaeophytes showed the strongest decrease across the whole study period (approx. -21.6%). While losses were strongest between the first and second period, changes between the second (1988 -1996) and third period (1997 – 2017) were marginally insignificant (i.e. their 95% CIs overlapped with 0, their 90% CIs didn’t). There was a tendency of mean occurrence in archaeophytes to further decrease also in the last observation period. In contrast, neophytes showed significant increases in occurrence of approx. +23.3% over the whole study period, with strongest increases between the second and third period. While the increase of neophytes compensated for the losses in natives and archaeophytes from the second to the third period, overall, the increase in neophytes did not level-off the decreases across the full study period (Table 1).

### 3.3. Changes in mean grid-cell species-richness

We detected significant changes in the estimates of mean grid-cell species-richness in all floristic status groups as well as across all species (Figure 3). While natives and archaeophytes showed consistent declines in mean grid-cell species-richness over the whole study period, neophyte species-richness showed only slight increases between periods two and three (i.e. 1988 – 1996 vs. 1997 – 2017). Losses were strongest for archaeophytes (−19.3%). As for changes in mean nationwide occurrence, increases in mean grid-cell neophyte richness did not counterbalance the decreases in the other floristic status groups, causing a net decrease of approx. -1.9% in mean grid-cell species-richness per decade across all species. Overall, the species-richness trends were similar to mean species occurrence trends.

**Figure 3:**
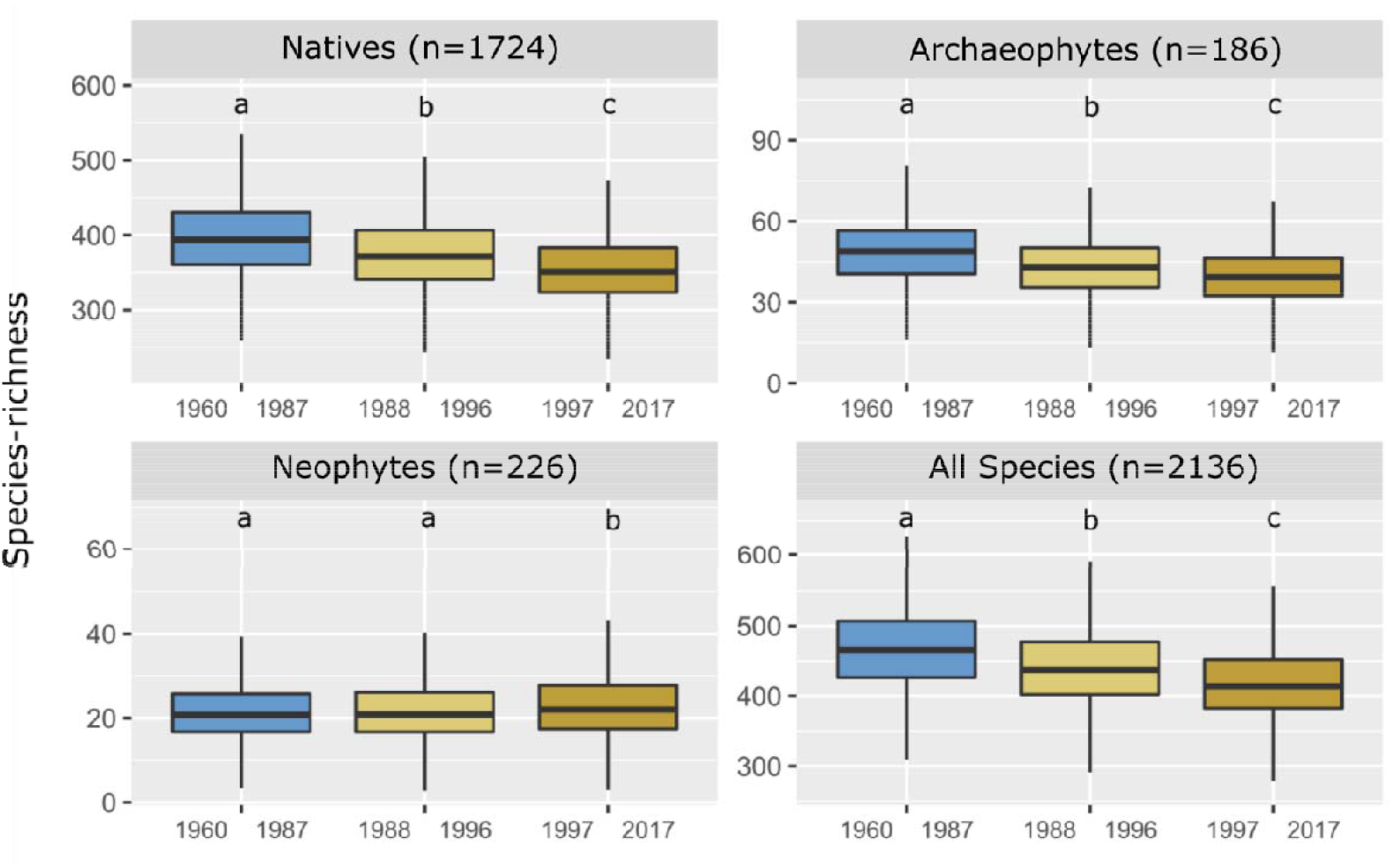
Changes in German-wide mean grid-cell species-richness (SOP_Grid_) across the three periods. Numbers in subplot headers indicate numbers of species included in the respective floristic status group. Letters indicate significant differences based on the temporal component in Model 2, accounting for spatio-temporal dependencies in the data (c.f. main text).

### 3.4. Spatial patterns of species-richness change

While the relative changes in grid-cell species-richness of archaeophytes and neophytes showed strong spatial heterogeneity, changes in native species-richness were more uniform, with consistent declines in all grid-cells (Figure 4; Electronic appendix, Table S4). Archaeophytes were also consistently declining but the magnitude of the decline was more variable, reaching the largest declines out of all floristic status groups in some grid-cells. Neophyte changes were most spatially variable and included regions of both decrease (especially in the north-east) and increase (especially in some southern regions). As for changes in occurrence, changes in species-richness of archaeophytes and natives were strongest from the first to the second period, whereas for neophytes they were strongest between the second and third. The spatial patterns of changes across all species closely followed those of natives (the most speciose floristic status group). Therefore, we will omit maps on the spatial patterns across all species from the following. For species-richness estimates and absolute changes see Electronic appendix (Figures S3 and S4, respectively; see also interactive maps: https://shiny.idiv.de/de25geka/PCHM/). While losses in archaeophyte species-richness from the first to the second study period were almost evenly spread throughout the country, there are regions with lower changes in archaeophyte diversity between the second and third period (Figure 4).

**Figure 4:**
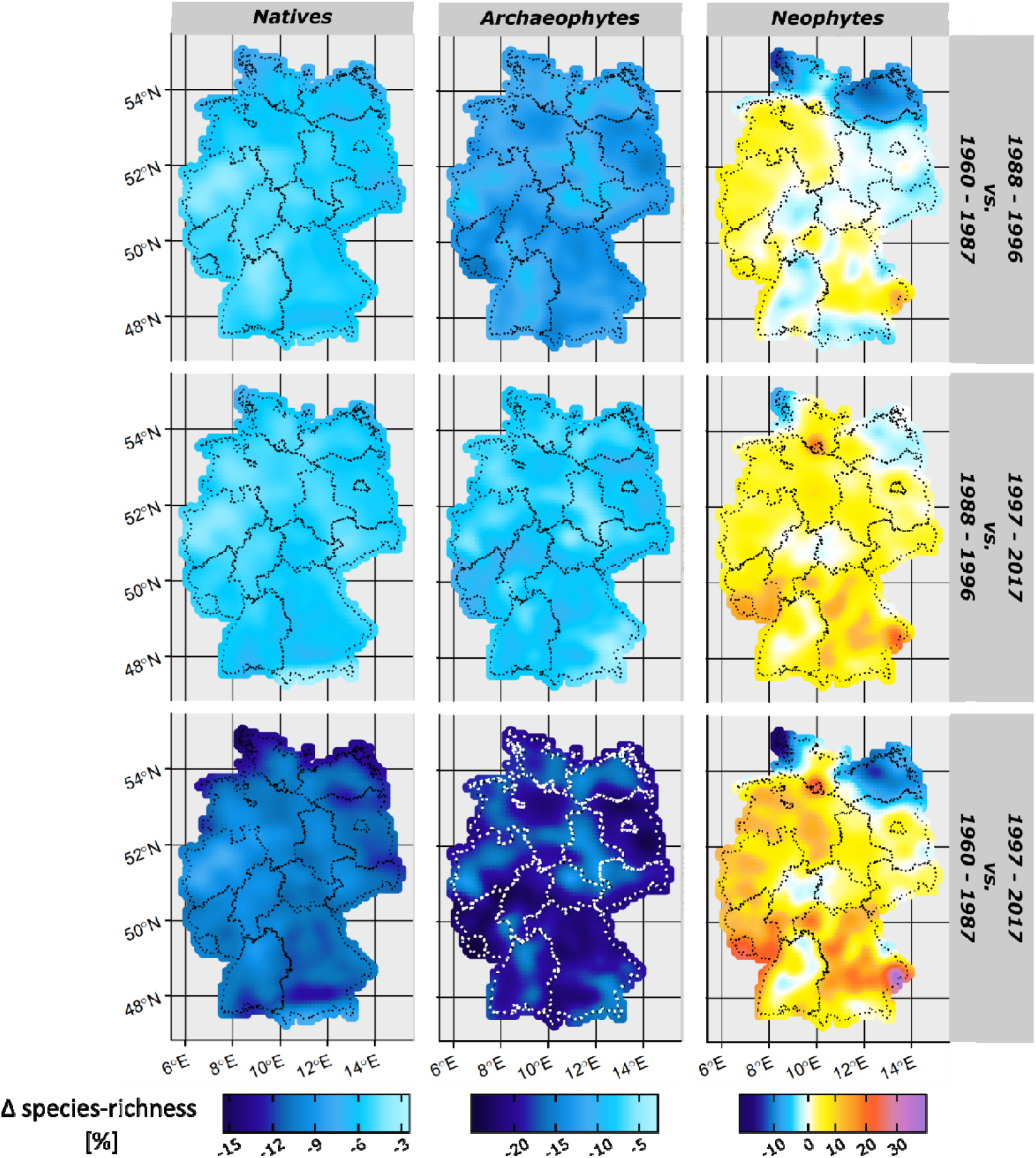
Relative changes in grid-cell species-richness and their spatial variability across the three study periods. Decreases are shown in blue, increases in yellow to purple. Dotted lines demark federal state boundaries.

Hotspots of changes in species-richness across the full study period are shown in Figure 5. Since we did not detect any grid-cells with increases in species-richness for natives and archaeophytes, hotspots of increases are only shown for neophytes. We were able to identify ten distinct regions (Figure 5, regions numbered from north to south), of which some are spatially congruent for two or even all three floristic status groups, while direction and magnitude may differ in space and/or time. The overlap of grid cells in hotspots of change was low for natives vs. archaeophytes (4.3%) and archaeophytes vs. neophytes (8.5%), but higher for natives vs. neophytes (24.6%, mostly in the coastal regions, Regions 1 and 3, Figure 5). Directions and strength in hotspots of species-richness change were not spatially correlated across the floristic status groups (F= 0.69, DF= 2 and 0.71, p= 0.68). We found strong declines in species-richness of natives and neophytes along the coast (Regions 1 and 3). Native species-richness also declined in the far east of Germany (Region 5) and the south (Regions 9 and 10). Regions of the strongest declines in archaeophytes were found in the southwestern parts of Germany (Region 6, but also 7 and 9) and in the northwest (Regions 3 and 4). Neophytes increased particularly strong in the southern half of Germany (Regions 6 – 8), but also in more northern parts (around Hamburg, Region 2). Regions with losses in natives did not match with regions gains in neophytes; similarly, hotspots of archaeophyte losses to not necessarily match those of neophyte gains (except regions 6 and 7).

**Figure 5:**
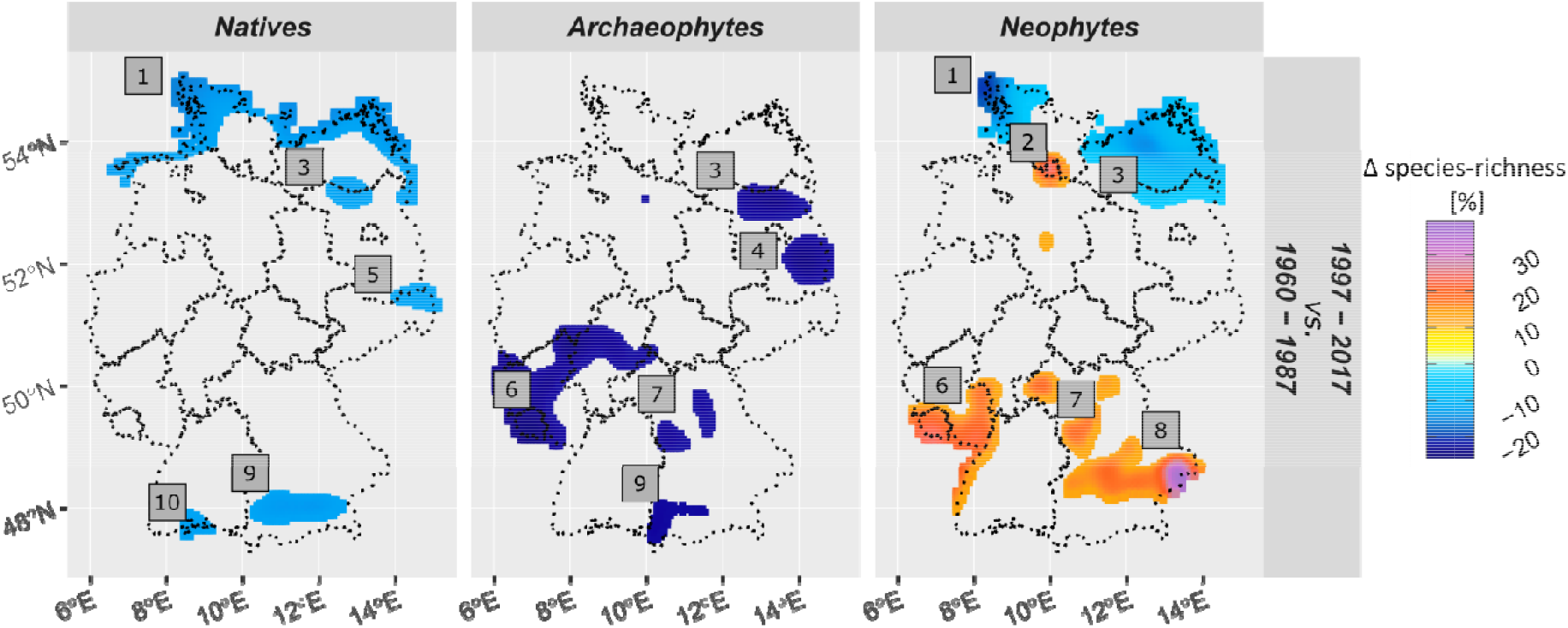
Hotspots of species-richness change between 1960-1987 vs. 1997-2017. Hotspots are defined as the lower and – in the case of neophytes – upper 10^th^ percentile of changes. Numbers depict geographical regions that can be roughly identified as (1) Schleswig-Holsteinische Geest and Wadden Sea; (2) Hamburg and Elbe Estuary; (3) Mecklenburg Large Lake District; (4) Liberose terminal moraine region; (5) Upper Lusatia; (6) Saarland and Hunsrück/Eifel region; (7) Main-Spessart and Middle Franconia; (8) Unterbayerisches Hügelland; (9) Alpine foothills and (10) Lake Constance Region; Dotted lines demark the state boundaries. For interactive maps see https://shiny.idiv.de/de25geka/PCHM/.

## 4. Discussion

Based on the collation of the largest databases on plant occurrence records in Germany to date, and correcting for reporting bias, we found significant declines in the German-wide occurrence of approx. 71% of all investigated vascular plant species over the last six decades. A much lower proportion (29%) of species showed increases in nation-wide occurrence probabilities. The increases in neophyte occurrence and species-richness did not compensate for nation-wide losses of other species, which led to a significant decrease of approx. -2% in mean species occurrence as well as mean grid-cell species-richness in Germany over the last six decades; Temporal and spatio-temporal dynamics differed between floristic status groups. We provide evidence that the majority of plant species in Germany shows a decline in occurrence and that decreases in plant species diversity are widespread across most regions throughout Germany.

The present study overcomes some of the main critiques on large-scale studies of biodiversity change summarized by Cardinale et al. (2018). The spatial representation of our data is not restricted to certain plots or locations in Germany that may cause a spatial bias towards certain regions or habitat types. In fact, our data covers all 5 × 5km German grid-cells. Moreover, since our analysis does not treat datasets independently but as an amalgam of different sources, underlying biases in the potential drivers of certain datasets (e.g. some vegetation relevés may have originated from success control of restoration projects) are reduced by combining these data with e.g. grid mapping for atlases that do not include such biases. Moreover, our study differentiates between changes in natives vs. non-natives.

### 4.1. Species-specific changes in occurrence probability

With approx. 56% of all vascular plant species in Germany (2136 out of 3868 taxa, raised to the harmonized taxonomy; the complete flora of Germany comprises a total of 5256 taxa, including subspecies and variants, Metzing et al. 2018), our study covers a major part of the German flora. Most of the species omitted from this study occurred only in a single time step or had less than 23 observations (c.f. Table S2). Thus, our analysis included all of the moderately common to common species, except very few *Oenothera, Rubus* and *Taraxacum* species, due to unstable taxonomical concepts. The study did not comprise very rare plant species, those that have gone extinct before 1987 or those that entered the German flora after 1996. A study by Lennon et al. (2004) demonstrated that, although rare species can constitute a major part of the species pool in a region, spatial structures of species-richness are typically dominated by the more common species. Therefore, the detected temporal and spatio-temporal patterns can be assumed to be robust estimates for change in species-richness. Moreover, our approach is rather conservative in terms of species presence: a species that is absent from a grid-cell in our original dataset, but detected (even only once) in the neighborhood of the respective grid-cell causes the occurrence probability in the respective grid-cell to be greater than zero, instead of it being rated as absent. However, although we did everything to ameliorate biases due to differences in local recording effort in our data and accounted for spatio-temporal biases, we cannot rule out the possibility that our data still contains some artifacts. Nonetheless, our work represents a major advance in the investigation of national plant diversity, and complements existing German structured monitoring schemes that mainly focus on rare, threatened or invasive species (Mitschke et al., 2005). However, we emphasize that close inspection of individual species trends and discussions with experts is crucial before making inferences. Moreover, the species-specific results provided in our analyses should be interpreted considering the spatial scale of the grid-cell level. On the grid-cell level, a species can only be rated as absent after the last individual in a grid-cell has gone extinct (Chase et al., 2019). Thus, the results of our analyses refer to occurrence, and not abundance. For example, the species with the strongest decline of approx. -99.9% (*Anagallis tenella*), indicating a near-extinction, has been recognized as “threatened with extinction” (RL 1) in the German Red List of 2009 (BfN, 2009) but has been changed to “endangered” (RL 2) in 2018 (Metzing et al., 2018). While there are only three remnant occurrences of this plant species (shown in Figure 2) that have been decreasing in size in the last decades, populations within these remnants have stabilized due to nature protection measures (Metzing et al., 2018; Raabe et al., 2012). By contrast, the species with the most extreme increase, *Senecio inaequidens*, has been recognized to be expanding since the early 1960s (Griese 1996), mainly along the railway and road network. Our approach can identify such large-scale changes, but it cannot replace specialized, local-scale investigations such as population-based studies, e.g. for red list assessments.

An increased number of endangered plant species in Germany has recently been reported in the German Red List of Endangered Plants (Metzing et al., 2018). Our results report an even higher total number of species in decline - many of them common species - which should raise the attention of decision makers and the general public. Like in the present study, a study in north-east Germany reported that approx. 60% of the 355 studied species were declining and that moderately common species declined strongest (Jansen et al., 2020). Similarly, in north-west Germany Bruelheide et al. (2020) reported significant declines for a large number of plant species, mainly herbs.

### 4.2. Changes in occurrences of floristic status groups and their spatial patterns

The general loss in species-richness across all species was dominated by the declines among native species, which is the most speciose group. We were able to show that the patterns of change do not only vary according to floristic status, but also in spatial and temporal patterns. An investigation of the causal relationships between large-scale measures and potential drivers was not in the scope of the present study. However, we can discuss the spatial patterns, especially the hotspots of change, with respect to knowledge from local, small-scale studies. For example, in the German coastline regions (also Region 1 in Figure 5), a number of coastal macrophytes, e.g. *× Calammophila baltica* (Schrad.) Brand. and *Leymus arenarius* (L.) Hochst., two coastal grass species, were predicted to decline in climate envelope models for the German coastal regions due to climate warming (Metzing, 2010). Likewise, Kastler & Michaelis (1999) and Eggert et al. (2006) reported that *Zostera marina* L. and *Z. noltei* Hornem., two submerse seagrasses, are declining due to increased sea temperatures, salinity and eutrophication in the wadden sea. A study on the scale of the northeastern federal state of Mecklenburg-Western Pomerania by Jansen et al. (2020) found also other typical coastal species such as *Salsola kali L.* or *Triglochin maritimum L.* declining between 1980 and 2000, probably due to hampered coastal dynamics and reduced grazing of coastal grasslands. All mentioned species show declining in occurrence also in our analysis, indicating that the findings of the more local studies hold also on larger scales.

The decline in the occurrence of archaeophytes shown in our study, has also been reported from small-scale studies, and was often explained by an increase in land-use intensification (Baessler & Klotz, 2006; Comin & Poldini, 2009; Knapp et al., 2010; Leuschner et al., 2013; Meyer et al., 2015). A study by Baessler & Klotz (2006) demonstrated, that arable weeds decreased in intensively used arable fields, whereas opportunistic ruderal species strongly increased in species-richness. Our data does not allow to assess causal relations between agricultural land use and biodiversity change. For future research it may therefore be worthwhile to investigate these connections in more detail.

Anthropogenic influences have been shown to be not always detrimental for local biodiversity in general: archaeophytes arrived in Central Europe only because of human influence, neophytes have been reported to profit from urbanization (Knapp et al., 2010; Kühn & Klotz, 2006) and human trade and transport (Rejmánek et al., 2005). In addition, several studies have shown that estuaries may especially be prone to the establishment of non-native species for various reasons (e.g. Wolff, 1999). For example, as demonstrated for the Elbe estuary, rivers may sediment excess nutrients there and the brackish water habitats often show the greatest ‘indigenous species minimum’, so that more alien species can potentially establish (Nehring, 2006). In support for this, we detected strong increases in neophyte species-richness around the city of Hamburg and the Elbe estuary (Region 2), which has the biggest German harbor, with strong international trade.

The area with strongest increases in neophyte species-richness in the present study (Region 8, Figure 6), has been reported as a region of strong increases in the establishment and expansion of neophytes from the climatically mild Danube plains in a local-scale study by Sompek et al. (2017). The authors claim that this increased expansion is due to an increase in habitat suitability caused by climate change. Neophytes are known to rapidly colonize newly available habitats, using mild valley refuges as a starting point for expansion (Rejmánek et al., 2005). Our spatial maps of changes in grid-cell species-richness confirm this spread and also show that the expansion of neophytes is widespread. Similarly, a study in the nature reserve in the northern part of the upper Rhine valley (close to Region 6) conducted by Vor & Schmidt (2008) reported an increase in neophyte species-richness compared to atlas data from 1993 (Lang & Wolff, 1993). Based on our maps, we can show that this increase in neophytes is more widespread than the upper Rhine valley alone.

While acknowledging that our study cannot give empirical evidence for causal relationships, we conclude that the spatio-temporal patterns of change in the national plant biodiversity are highly variable which is evidence for a complex interplay of drivers of biodiversity change. As demonstrated by the comparatively low level of spatial congruence in the detected hotspots of species-richness and the lack of correlation between the hotspots of species-richness change among the different floristic status groups, these factors act locally, and affect different species in different ways. Nevertheless, the fact that on the national level, net plant species-richness is declining so predominantly and permanent is alarming. While our study is focused on Germany, we have no reason to believe that these changes are only limited to this country. Plants, as primary producers, play pivotal roles in ecosystems and changes in their biodiversity may cascade throughout the food web (Jansen et al., 2020) and influence ecosystem functioning across trophic levels (Emmerson et al., 2016; Schuldt et al., 2018). For example, changes in the floristic composition of habitats in north Germany have been shown to result in lower nectar availability, with probably negative effects on pollinating insects (Bruelheide et al., 2020). It is therefore possible that the detected large-scale changes in plant biodiversity is connected to recent insect declines (Hallmann et al., 2017; Seibold et al., 2019).

## 5. Conclusion

Declines in vascular plant biodiversity are widespread in Germany and are not restricted to taxa listed as being endangered in national Red Lists, but is apparent in more than 70% of plant species studied here, and thus in 40% of all moderately common to common vascular plant species in Germany. Urgent action is needed to halt this biodiversity loss. Our approach demonstrates how existing large datasets can be combined and used for robust trend analysis. Existing data should also be collated from other states in Europe and globally. The data integration and analysis approach used in this study is comprehensive and robust to different methodological biases. It can be applied to other large-scale research using heterogeneous occurrence data, given it can be harmonized to a common fine-scale grid. This makes our approach valuable for other projects, such as the growing Living Atlas community (Brenton et al. 2018) and may also help to inform and create new, more collaborative monitoring schemes that integrate knowledge and data from different actors in nature protection (e.g. governmental, academical and volunteer). Such new schemes should also include long-term monitoring of common species. Clearly this is an ambitious endeavor, which can only be accomplished through joint efforts of a variety of stakeholders and should be underpinned with the financial and legislative power of (inter)national institutions. However, such approaches must not question the need for monitoring projects, which are still necessary and have the potential to identify the drivers of biodiversity change.

## Supporting information

Technical details

## 6. Acknowledgments

We are grateful to the many individuals that were involved in gathering the vast amount of data combined in our dataset and to those who helped to mobilize the data, especially to those that contributed to large data-collections like FlorKart and the vegetation databases. Further, we are grateful to C. Storm, T. Heinken, T. Dittmann, V. Wagner, I. Dörfler, J. Reinecke, S. Kühn, T. Naaf, J. Kolk and M. Wulf for the kind contribution of relevé data. We also thank Karsten Wesche for the valuable discussion on the most extreme species-specific changes. The present study is an outcome of the project sMon of the German Centre for Integrative Biodiversity Research (iDiv) Halle-Jena-Leipzig, funded by the German Research Foundation (DFG - FZT 118, 202548816). Time and effort were supported by sDiv, the Synthesis Centre of iDiv. sMon is a collaborative project between iDiv and the German Federal Agency for Nature Conservation (Bundesamt für Naturschutz, *BfN*), conservation agencies of the 16 federal states, natural history societies, natural history museums, taxonomic experts and scientific institutes.

