## Supporting Figures for "Widespread decline in plant diversity across six decades"

**Electronic appendix to Eichenberg et al. (2020)**

**Supporting Figures**

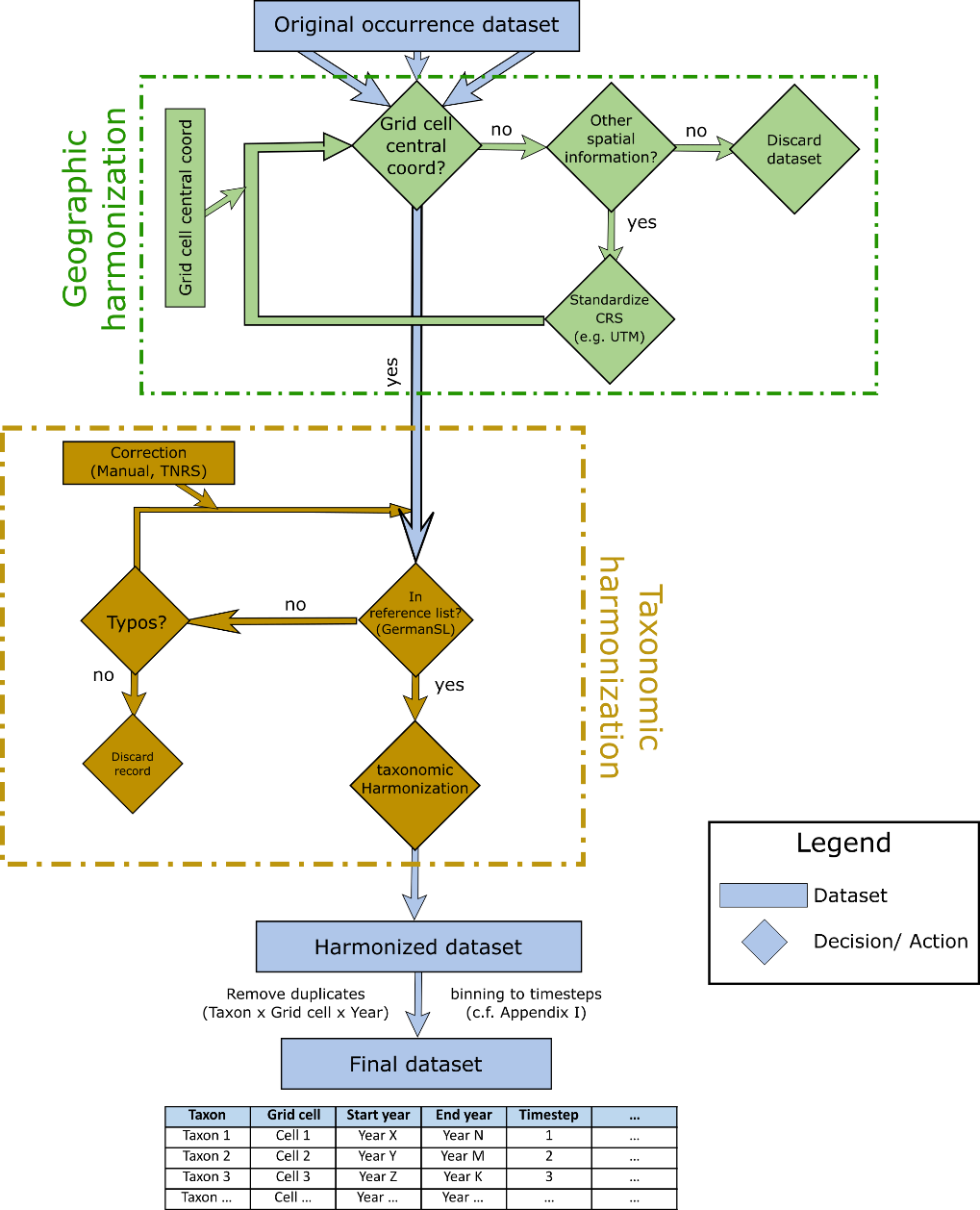

Figure S1: Flowchart for the workflow in geographic and taxonomic harmonization. Any potential dataset on occurrence records should be first standardized to a common coordinate reference system (CRS), preferably in a format that allows a straightforward calculation of distances in km or m. Taxonomic reference list can be any accepted list (for Germany here German SL, Jansen & Dengler 2008); UTM: universal Transverse Mercator; TNRS: taxonomic name resolution service.

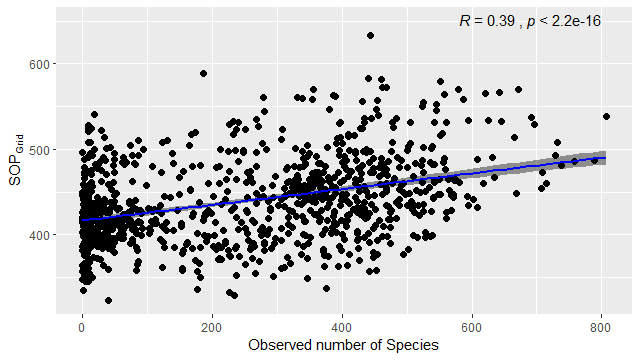

Figure S2: Correlation between predicted species-richness (SOP_Grid_) and Observed species-richness in grid-cells that can be assumed to be well sampled according to Kühn et al. 2006 (i.e. based on the presence of 50 ubiquitous in the FlorKart dataset; see Table S1). R= Spearman correlation coefficient.

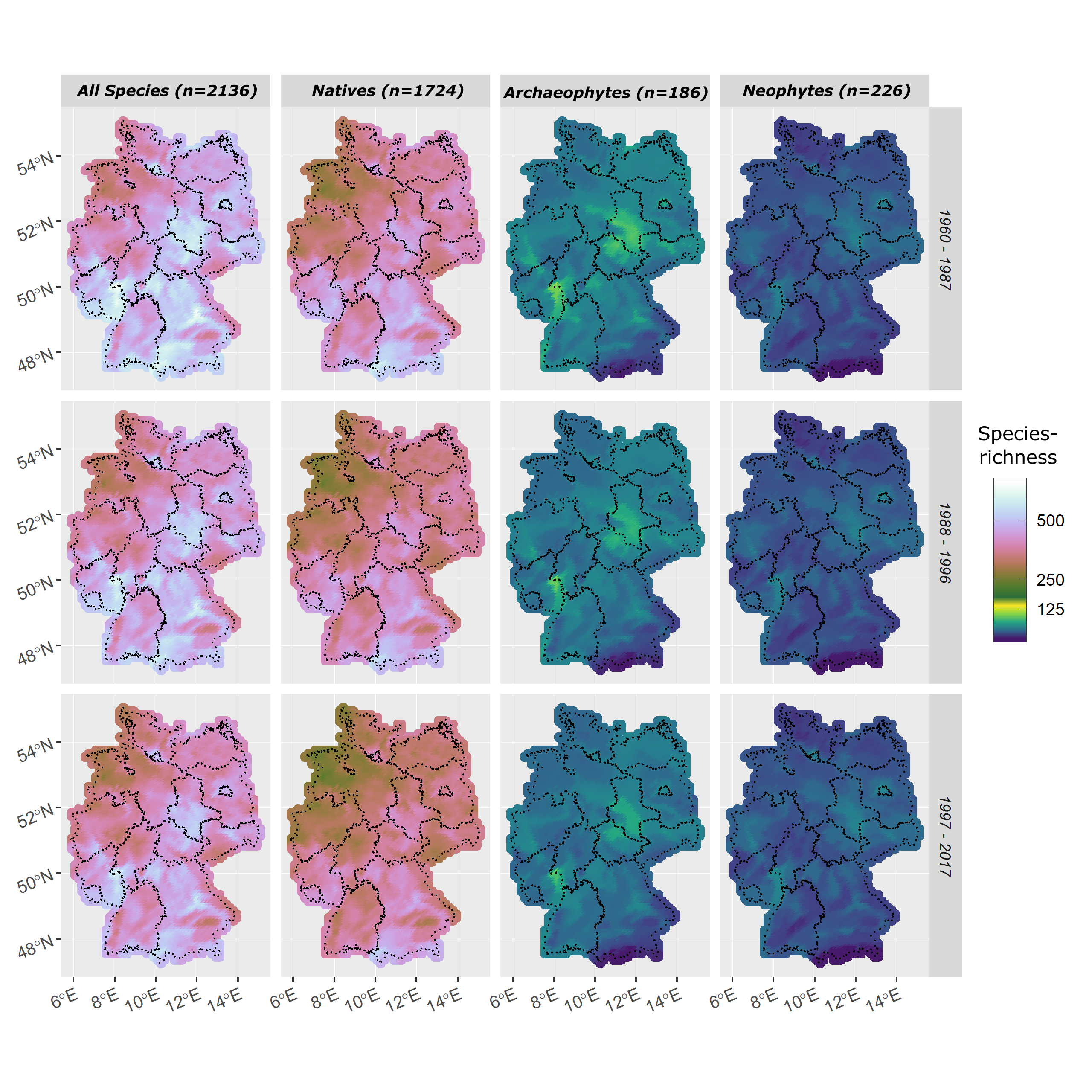

Figure S3: Spatial distribution in species-richness across Germany for the three study periods. Species-richness is calculated as the sum of occurrence probabilities of all species in a grid-cell.

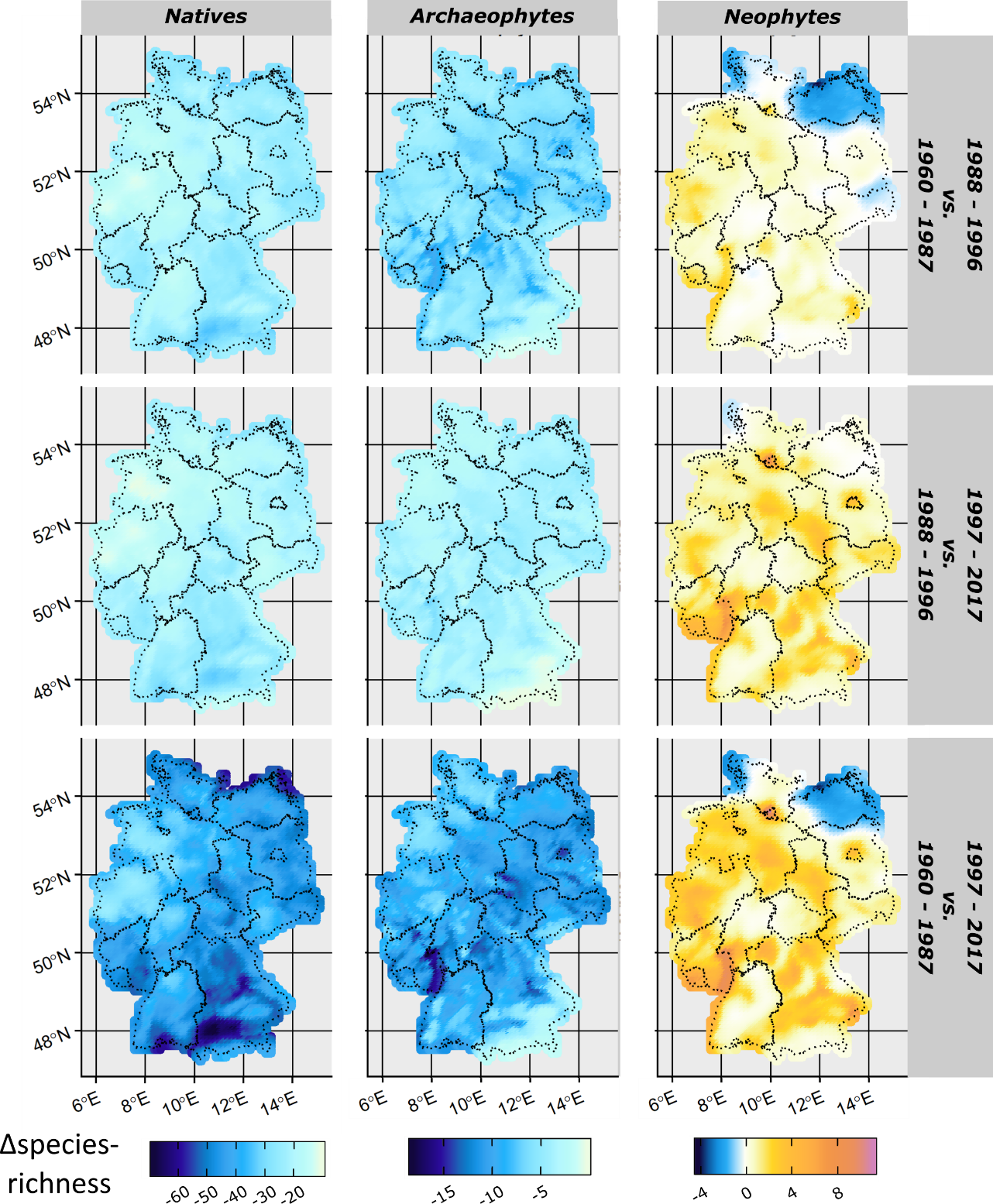

Figure S4: Absolute changes in grid-cell species-richness and its spatial variability across the three study periods. Decreases are shown in blue, increases in yellow to purple.
