## Supplemental Table 1 for "Widespread decline in plant diversity across six decades"

**Electronic appendix to Eichenberg et al. (2020)**

Supporting Tables:

Table S1: Data sources, type of data (occurrence/relevé) and geographic properties as well as number of occurrence records and temporal extent of the data compiled in this study. MTB_Q= German Raster Quadrant (MesstischBlattQuadrant, i.e. approx. 5 x 5 km)

| **Data source** | **Type** | **Georef** | **n** | **Start Year** | **End Year** |
| --- | --- | --- | --- | --- | --- |
| FlorKart | Occurrence | MTB_Q | 24169633 | 1960 | 2013 |
| Landesamt für Umwelt und Geologie, Thüringen | Occurrence | Gauss-Krüger | 879 | 1966 | 2016 |
| Landesamt für Umwelt, Bayern | Occurrence | Gauss-Krüger | 884579 | 1961 | 2017 |
| Flora Mecklenburg-Vorpommern | Occurrence | MTB_Q | 653788 | 1960 | 2017 |
| BfUE Hamburg | Occurrence | DK25, Gauss-Krüger | 53514 | 1999 | 2017 |
| Landesamt für Umwelt, Sachsen-Anhalt | Occurrence | Gauss-Krüger | 355 | 1990 | 2012 |
| Landesamt für Landwirtschaft, Bayern | Occurrence | Gauss-Krüger | 88455 | 2002 | 2013 |
| LLUR, Schleswig-Holstein | Occurrence | Gauss-Krüger | 96955 | 1963 | 2017 |
| vegetweb | Relevé | Mixed | 1953277 | 1961 | 2014 |
| GVRD | Relevé | Mixed | 877173 | 1960 | 2015 |
| Institute of Biogeography, Univ Bayreuth | Relevé | Gauss-Krüger | 6066 | 1989 | 2013 |
| Biodiversity Exploratories | Relevé | WGS84 | 5014 | 2008 | 2009 |
| Private (Storm, C., Heinken T., Dittmann, T., Wagner, V., Dörfler, I., Reinecke, J., Kühn, S.L., Naaf, T., Kolk, J., Wulf, M.) | Relevé | Mixed | 13708 | 1960 | 2016 |
| Storm (Uni Darmstadt) | Relevé | Gauss-Krüger | 9417 | 1995 | 2016 |
| Leutratal & Cospoth, Univ. Jena | Relevé | Gauss-Krüger | 126386 | 1972 | 2016 |
| Sum (Above) |  |  | 28939199 | 1960 | 2017 |
