## Supplemental Table 4 for "Widespread decline in plant diversity across six decades"

**Electronic appendix to Eichenberg et al. (2020)**

Supporting Tables:

Table S4: Mean, minimum, maximum and standard deviation of % changes in German-wide grid-cell species-richness across the three time periods. For a spatial representation see Figure 5.

| Status | Period | Minimum | Mean | Maximum | sd |
| --- | --- | --- | --- | --- | --- |
| Natives (n=1724) | 1960 - 1987  vs.  1988 - 1996 | -8.16 | -5.55 | -3.69 | 0.75 |
|  | 1988 - 1996  vs.  1997 - 2017 | -7.56 | -5.39 | -2.8 | 0.64 |
|  | 1960 - 1987  vs.  1997 - 2017 | -15.1 | -10.64 | -7.41 | 1.12 |
| Archaeophytes (n=186) | 1960 - 1987  vs.  1988 - 1996 | -16.06 | -11.94 | -7.66 | 1.49 |
|  | 1988 - 1996  vs.  1997 - 2017 | -11.04 | -7.96 | -3.4 | 1.2 |
|  | 1960 - 1987  vs.  1997 - 2017 | -24.51 | -18.94 | -12.59 | 2.25 |
| Neophytes (n=226) | 1960 - 1987  vs.  1988 - 1996 | -13.66 | 0.72 | 13.8 | 4.19 |
|  | 1988 - 1996  vs.  1997 - 2017 | -6.97 | 5.33 | 21 | 3.63 |
|  | 1960 - 1987  vs.  1997 - 2017 | -19.58 | 6.19 | 37.7 | 7.31 |
| All Species (n=2136) | 1960 - 1987  vs.  1988 - 1996 | -8.74 | -5.89 | -3.75 | 0.79 |
|  | 1988 - 1996  vs.  1997 - 2017 | -7.63 | -5.09 | -2.69 | 0.72 |
|  | 1960 - 1987  vs.  1997 - 2017 | -15.7 | -10.68 | -6.98 | 1.26 |
