## Supplementary material for "Widespread decline in plant diversity across six decades": Technical details

**Electronic Appendix to Eichenberg et al. (2020)**

**1. Technical details I: Frescalo Analyses**

1.1. **Defining the study periods**

The FRESCALO algorithm is distinctly designed to evaluate data on species occurrences in grid cells, which are often collected by biological recording schemes, typically with the intention of publishing an atlas (Hill 2012). Such data is heterogeneous in its nature, in the sense that it may suffer from varying efforts on species recording, depending on the respective recorder, regions under study, taxa, or timeframes included. Using a moving window in space that, in each iteration, has a focus grid-cell centered within a neighborhood of adjacent grid-cells, the algorithm uses the relative frequency of species occurring in the surrounding grid-cells to fill in species which are missing in the focus grid-cell but are likely to be there (although e.g. missed by the recorder). The neighboring gird-cells are referred to as “neighborhoods”. Neighborhoods size is defined by the user, and typically comprise 50-100 of the ecologically most similar grid-cells in the surrounding of a focus cell. “Ecological similarity” is a user-defined property and should comprise ecologically meaningful variables (e.g. climate, topography or other meaningful variables; for examples see section 3 below or Hill, 2012). In the study of Eichenberg et al., (2020), we defined a neighborhood size of 100 grid-cells, from which the algorithm took the 50 ecologically most similar ones into account. Being based on neighborhoods, the algorithm itself is robust to a certain amount of totally unsampled grid-cells (c.f. Hill 2012), as long as the missing grid cells are not too many within a neighborhood and ideally are missing at random in time and space.

The occurrence records from Germany, used in the study by Eichenberg et al. (2020, main text), are peculiar in the way that they are mainly collected by federal states. These federal states coordinate their efforts separately and do not always collect (or submit) their data at the same times. Therefore, it may happen that, while a majority of the country may have recorded data in one period, one or several federal states did not conduct recording projects in this period. Therefore, a federal state will not have data in this respective period; in these cases, the spatial gaps in the data are definitely not missing at random and the areas not covered by the data may be too big for the algorithm to fill in with sufficient accuracy.

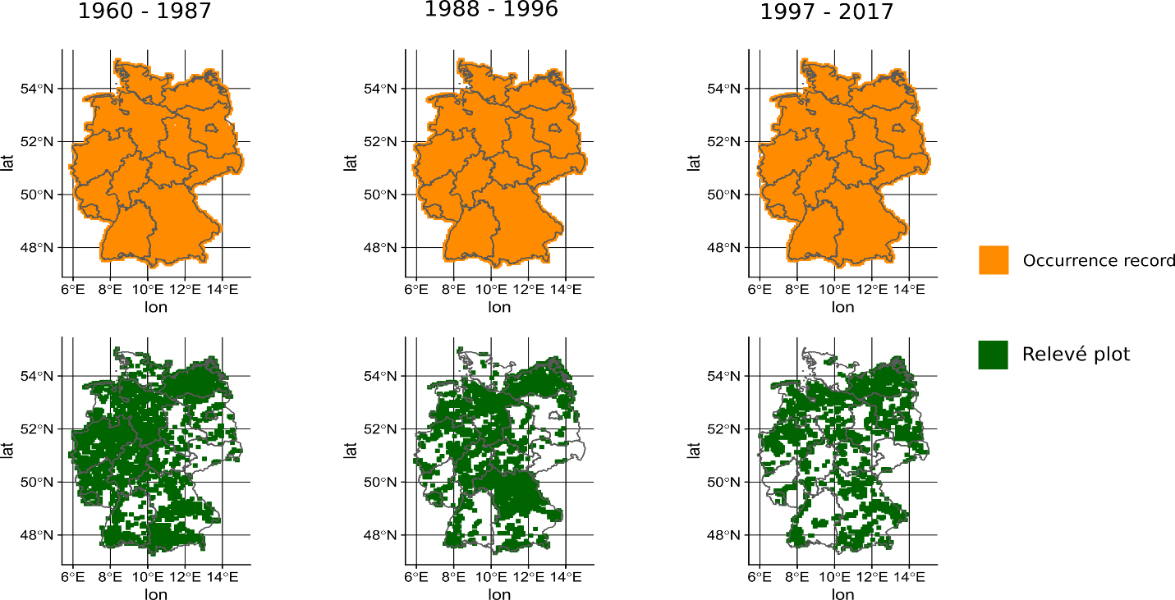

Figure A1: Spatial distribution of occurrence (top) and relevé (bottom) data across the defined study periods. In our conservative approach, we made sure that the occurrence record data covered the full area of Germany in every period. The community data from vegetation relevés was used to “boost” taxonomic information in the neighborhoods defined in the FRESCALO algorithm (c.f. Chapter 3 below).

Thus, the first challenge in an analysis using the FRESCALO algorithm is to define suitable periods that are wide enough to not show big spatial gaps that are not missing at random. Moreover, the periods should be chosen in such a way that the “taxonomic intensity” (i.e. the taxonomic focus of single recording projects) are not too narrow. An example here would be a project is especially focused only on a narrow taxonomic scale (e.g. only critically endangered species or species from a single genus, e.g. Rubus; only species of a single taxonomic or functional group such as invasive species). This is to ensure that, even if single (focus) grid-cells may be poorly sampled, the surrounding neighborhood is sufficiently well sampled to fill represent the expected species assemblages within the focus grid-cell.

We tackled this challenge in several ways:

a) We included data from different sources, i.e. pure occurrence records from mapping projects and extensive community data from vegetation relevés. While the former may be taxonomically less complete across time and space, the latter are especially designed to capture the full community of the respective plot under study, but are often not replicated in time. While mapping projects cover large areas (in Germany the full 5 x 5 km grid-cell), vegetation relevés are much smaller (e.g. 1 – 10 m², e.g. Dengler, 2009).
b) We made sure that the study periods are wide enough, so that the data from mapping projects cover all grid-cells in Germany (Figure A1), and that the data collected in these mapping projects do not focus in on a narrow taxonomic range. The taxonomic range was defined as the number of plant families that were recorded by a mapping project (categorized into five classes, see below as well as Figure A2). Vegetation relevés were always defined as the highest intensity class (i.e. class 5).

With this, we made sure that there are no spatial gaps in each timestep, and that the taxonomic bias is not too big in the occurrence records data. We then used the taxonomically complete, small scale data from vegetation relevés to “boost” the taxonomic completeness in the neighborhood of the moving focus grid-cell.

Identifying the suitable study periods was an iterative approach. The fact that the different periods span different time frames (approx. 2 decades in the first and last, but only 8 years in the second period) has the drawback to prevent us from calculating trends on a fine temporal resolution (e.g. years), but is robust enough to calculate these changes on a decadal basis. It is, however, not a drawback to the application of the FRESCALO algorithm itself, as it is not affected by unevenly spaced periods (Hill, 2012), given the periods are carefully balanced in terms of spatial and taxonomic coverage (see above).

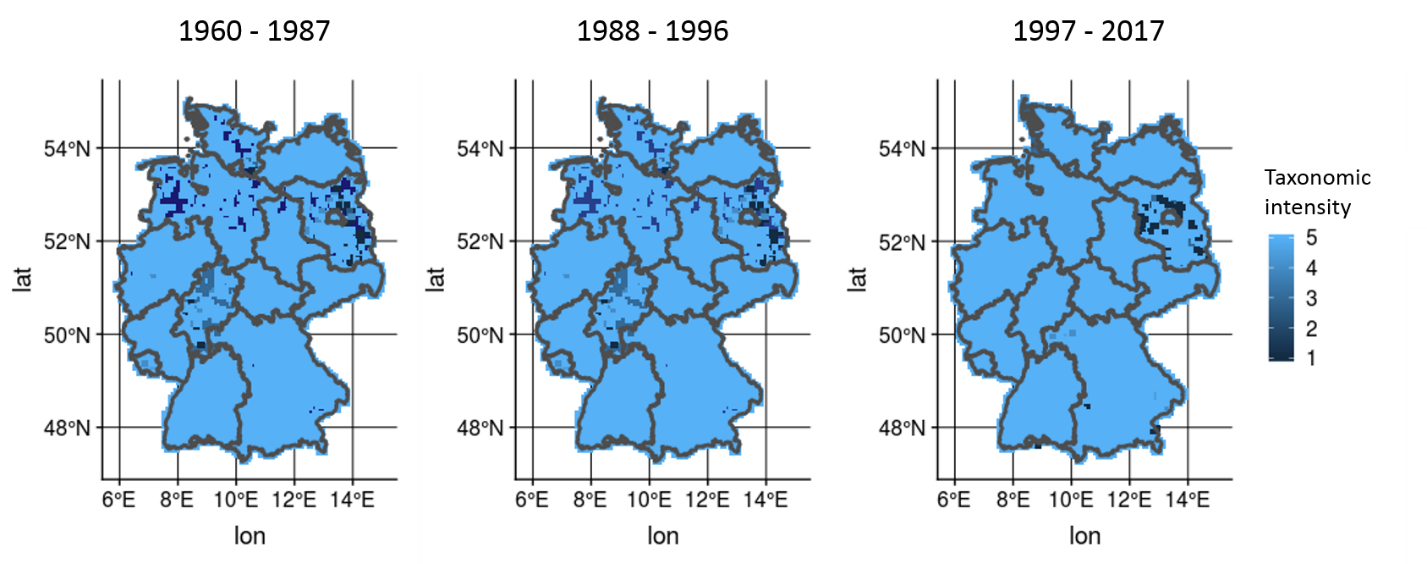

Figure A2: Spatiotemporal coverage of taxonomic range (i.e. median of number of recorded families in a grid-cell on a five-level scale per data source) in the respective study period. 1: 1 family; 2: 2 – 20 families; 3: 21 – 50 families; 4: 50 – 100 families; 5: >100 families. NOTE: Data from vegetation relevés were always treated as the highest intensity.

1.2. **Specifications of the FRESCALO algorithm**

The FRESCALO algorithm, as available in the R-package ‘sparta’ (version 0.1.48, August, 2015) requires two parameters to be specified by the user (Hill, 2012). The first parameter is the standard neighborhood frequency Φ, which reflects the expected frequency of species in the neighborhood. The optimal value for Φ can be computed automatically by the algorithm and was determined as 0.79 in Eichenberg et al. (2020, main text). The second parameter is the proportion of species within a neighborhood treated as benchmark species. The higher the proportion of benchmark species, the more species need to be found in a neighborhood for it to be considered as well sampled. In a preliminary study we tested different values of this parameter (10, 15, 20, 25%), but found no qualitative difference in the results (not shown). This is also in accordance with sensitivity test carried out by Hill (2012, supplementary material). In the main text, we therefore present the results obtained for a proportion of 20%. This amounted to approx. 70 – 320 benchmark species per neighborhood based on our raw data. Species that are rated as invasive in Germany (Nehring et al., 2013) were excluded from the list of possible benchmark species (see Bijlsma 2013).

1.3. **Defining the ecological similarities of neighborhoods**

In brief, neighborhood similarities for FRESCALO are calculated on the basis of spatial proximity (more distant neighbors have lower similarity) and other criteria, which can be user specified. For the Analyses in Eichenberg et al. (2020, main text), spatial proximity and ecological/topographical/climatic similarity were calculated as a product of the inverse spatial distance and similarity in 76 environmental variables (see below). So, closer and environmentally more similar sites get higher similarity values than close, but environmentally dissimilar neighboring sites. These values are then used to weight the local frequencies of species i in the respective neighborhood. With this information at hand, FRESCALO calculates the probability for species i being present in the target site j and period t in the neighborhood under study. For details on the calculation of neighborhood similarities, the interested reader is referred to Hill (2012) as well as the vignette of the R package ‘sparta’ (August 2015).

*Similarity variables:*

- Spatial distance: We calculated the central coordinates (harmonized to UTM Zone 32) of each raster quadrant in our dataset based on GIS information using the function st_centroid() function from the sf package in R (Pebesma 2018). The central coordinates were then used to calculate the geographic distances between the raster quadrants in each neighborhood as the Euclidean distance in km based on UTM coordinates.
- Environmental similarity: three measures of environmental similarity were chosen to reflect neighborhood similarity
  - topographic similarity: based on a 25m DEM freely available for the complete European Union (European Environment Agency 2017) we calculated the mean, maximum and minimum values for elevation, slope as well as the mean aspect of each raster Quadrant in Germany. In addition, we calculated the ratio of projected 2D to extrapolated 3D surface as a measure of surface roughness for each raster quadrant (Rashid 2010).
  - edaphic similarity: Edaphic similarity was calculated as the percentage of area of each raster quadrant covered by the 62 soil types defined in the “Bodenübersichtskarte 1:2000.000 (BÜK2000)” available from <https://www.bgr.bund.de/DE/Themen/Boden/Informationsgrundlagen/Bodenkundliche_Karten_Datenbanken/BUEK_2000_3000_5000/BUEK2000/buek2000_node.html>.
  - Climatic similarity: climatic similarity was calculated based on the climatic data available in a 1x1 km grid for Germany from 1902 - 2017 by the German Weather Service (Deutscher Wetterdienst, DWD) openly available under
    <ftp://ftp-cdc.dwd.de/pub/CDC/grids_germany/annual/>. We used values for yearly minimum, maximum and mean values of temperature as well as mean precipitation for characterizing climatic similarity. The data for each year were aggregated to the German ordinance grid (approx. 5 x 5 km) using the extract() function available from the R-package ‘raster’ (Hijmans 2019). Subsequently, we calculated the average of these across the study period (1980 - 2009) for each raster quadrant.

The R code to run the Frescalo Model is given in the gray boxes:

```
### START of Script

FrescaloResults<- frescalo(Data = FrescaloData,
         frespath = myFrescaloPath, # Path with Frescalo_3a_DE.exe
         time_periods = timeslices, # Dataframe with start and end years of timesteps
         non_benchmark_sp= nobenchm_specs, # List of species with unstable taxonomy and
                                             invasive species
         site_col = 'MTB_Q', # Column in FrecaloData with grid cells
         sp_col = 'Taxon', # Column in FrescaloData with Taxonomic IDs
         start_col = 'Start_Year', # Column in timeslices with Start Year
         end_col = 'End_year', # Column in timeslices with End Year
         Fres_weights = Fresweights, # neighborhood weights specified as described above
         sinkdir = c(“#YourOutputDirectory’”), # output directory (NOTE: Generic, here)
         phi=0.79, ## neighborhood frequency; optimal value determined in a preliminary study
         alpha=0.20) # percentage of species in neighborhood used as benchmark species

save(FrescaloResults,”#YourOutputDirectory/FrescaloResults.rda”)

### END of Script
```

All values were compiled in a single matrix with raster quadrants as rows and variables as columns. We calculated the Euclidean distances of all raster quadrants. These were then used as dissimilarities in the computation of neighborhood similarity via the createWeights() function in ‘sparta’. Here, we chose the 50 ecologically most similar grid cells out of a neighborhood size of 100 gird cells as relevant for the compilation of the list of benchmark species (see Hill, 2012).

1.4. **Detailed description and discussion of neighborhoods:**

Here we present a detailed discussion of four exemplary neighborhoods, based on a subset of the data extending across the German federal state Mecklenburg Western Pomerania. The neighborhood analysis reflected intuitive neighborhood similarity based on common sense and expert knowledge of the region (Figure A3). For example, the target site in neighborhood 1 (Figure A3, green dots) was located directly at the coastline of MV, close to the city of Boltenhagen. The grid-cells most similar to the target site are all located close to the coastline, whereas sites in the neighborhood that are situated more to the inside of the federal state (further away from the coast) are considered as less similar (i.e. they achieve lower weights). The similarity distribution of neighborhood 2 (purple) shows another feature of the similarity weights: even sites spatially localized closer to the target site may achieve lower weights, if they are environmentally less similar. The target site in neighborhood 2 is located inside the nature park “Nossentiner/Schwinzer Heide” (polygon not shown, but see <https://geodienste.bfn.de/schutzgebiete?lang=en>, CDDA-Code: 323290), a region that is rich in forests and features a high percentage of rivers and lakes. Sites within the neighborhood that also fall into this nature park are environmentally similar, thus they receive higher weights. On the other hand, the nature park ends at the east of the target site. Sites in the eastern part of the neighborhood are more dominated by an agricultural landscape with environmental conditions that strongly differ from the target site. The parameters that were used to calculate the neighborhood similarity do capture this difference very well, and consequently similarity weights for these sites are lower (i.e. dots are smaller). Then again, plots that lie more west in neighborhood 2, again are located in a nature park called “Dobbertiner Seenlandschaft” (<https://geodienste.bfn.de/schutzgebiete?lang=en>, CDDA-Code: 320378). The “Dobbertiner Seenlandschaft”, similar to the “Nossentiner/Schwinzer Heide” features moist and less intensively used landscapes, and consequently sites here are weighted as more similar to the target site.

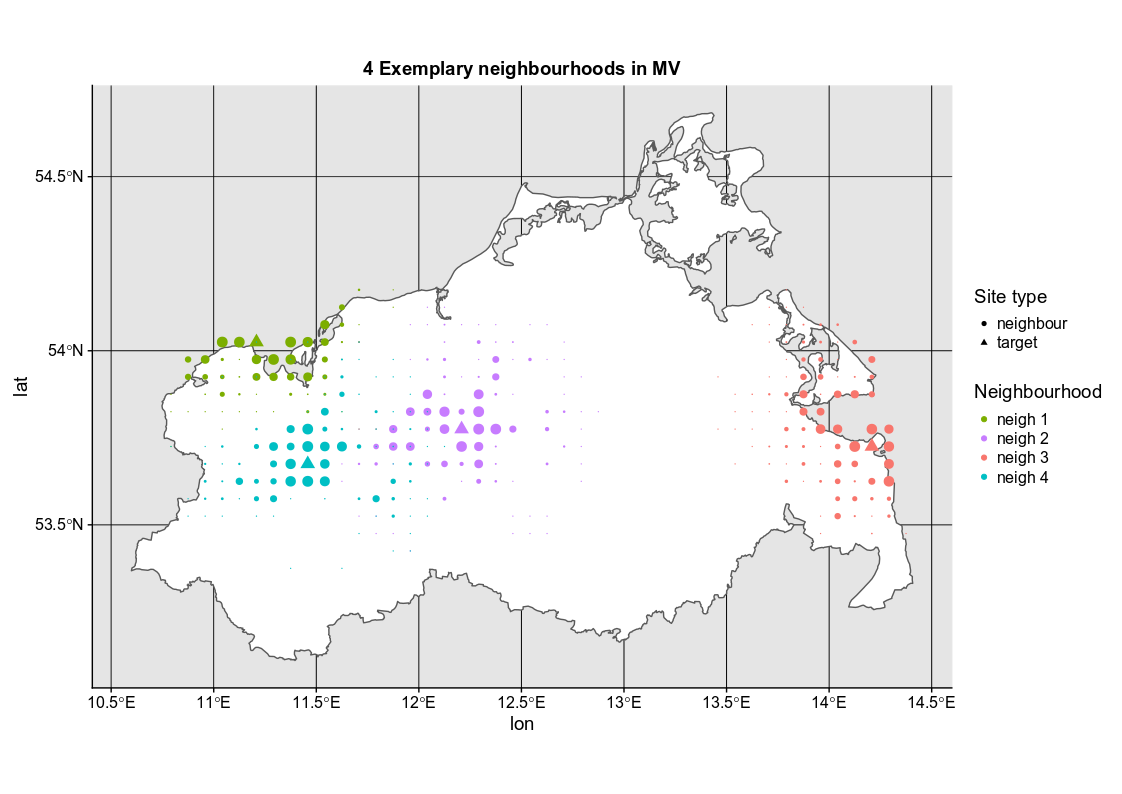

Figure A3: Four exemplary neighborhoods (neigh 1-4) and their corresponding weights (size of the symbols) in Mecklenburg-Vorpommern. The weights were calculated based on the spatial distance as well as information on edaphic, topographic and climatic similarities of the raster quadrants. Triangles: target site of the algorithm, circles: sites in the neighborhood. Size of the circles correspond to higher (bigger) or lower (smaller) weights.

Interestingly, there are two neighborhoods which have overlapping sites (neighborhood 2 and 4). Here, the target site of neighborhood 4 (light blue dots) is located close to the city of Schwerin, a more urban characterized landscape. As would be expected from common sense, the overlapping sites in the neighborhoods receive different weights for the different neighborhoods of the respective target sites.

This detailed discussion demonstrates, that the definition of ecological similarity in the study of Eichenberg et al. (2020, main text) draws a realistic picture of “true” ecological similarity, thus preventing e.g. species specialized on coastal habitats to be erroneously projected into the non-coastal habitats in the inner landscapes. This is e.g. also reflected in the maps accompanying this publication (e.g. *Blysmus rufus* (Huds.) Link, Figure A4).

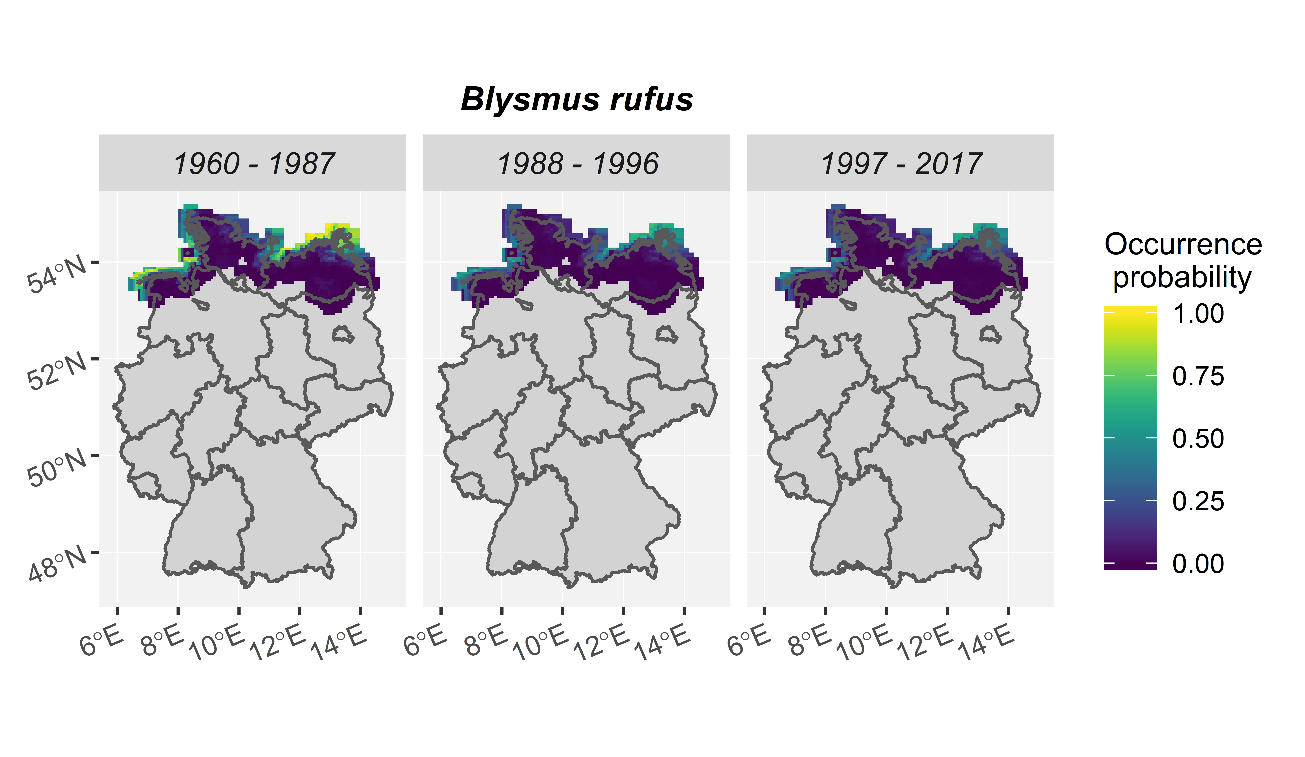

Figure A4: Occurrence probability of the maritime species Blysmus rufus (Huds.) Link. as calculated by the FRESCALO algorithm, based on the ecological similarities described above. This species is known to be mainly confined to salty, coastal habitats, with some inland populations in salty habitats.

1.5. **Code to produce spatial distribution maps from a Frescalo output object (according to the rationale in Bijlsma, 2013)**

**Note:** additional GIS Info is needed to add central coordinates to grid cells. For the current example these were taken from <https://geo.dianacht.de/mtb/mtbshape.zip>

Script will be continued on the next side

```
#### START of Script

library(sf)
library(ggplot2)
library(viridis)

MTB_Q<- #the GIS Data containing grid cell identifiers and central coordinates

## Extract number of grid cells
n_grid_cells<- length(unique(MTB_Q$MTB_Q))

load("./#YourFrescaloOutputDirectory/FrescaloResults.rda ")
MyData<- FrescaloResults
rm(FrescaloResults)

### 1. Disseminate Frescalo Object
### 1.1. Frequency information separately

frequencies<- MyData$freq

## housekeeping: some grid cells have only 4 digits, as the first digit in the origin is 0,  ## and thus are internally eliminated
frequencies$Location[nchar(as.character(frequencies$Location))==4]<- paste0("0", frequencies $Location[nchar(as.character(frequencies $Location))==4])
###### Creates an index to uniquely match Ops to obervations #########################
frequencies$index<- paste0(frequencies$Location,frequencies$Species)

### 1.2. Species IDs separately
species_ids<- MyData$trend$Species

## 1.3. Extract trendfactors (see Hill, 2012) for each timestep
n_timesteps= 3 ## define number of timesteps you investigated in the FRESCALO analysis

trendfactors<-

for (i in 1:n_timesteps){
data<- MyData$trend[MyData$trend$Time== unique(MyData$trend$Time)[i],
                    c("TFactor","Species","Time")]
data$index<- paste0(data$Species,data$Time)
assign(paste(“trendfacts_”,I, sep=””),data)
}
```

Script will be continued on the next side

```
### 2. Create a data frame
### 2.1. full size that can take Ops for species from the FRESCLAO output

for (i in 1:n_timesteps){
data<- data.frame(
           "MTB_Q"=as.character(rep(MTB_Q$MTB_Q,each=length(unique(species_ids)))),
           "Species"=as.character(rep(unique(species_ids),times=length(unique(MTB_Q$MTB_Q)))),
           "Timestep"=rep(i,times=length(unique(MTB_Q$MTB_Q))*length(unique(species_ids))),
           "lat"=rep(MTB_Q$lat, each= length(unique(species_ids))),
           "lon"=rep(MTB_Q$lon, each= length(unique(species_ids))))
data$prob<- NA
data$index<- paste0(data$MTB_Q,data$Species)
assign(paste(“spec_pres_t”, i, sep=””),data)
}

## 2.12 Shrinking the dataframe down to the actually populated grid cells (saves memory)

for (i in 1:n_timesteps){
data<- get(paste(“spec_pres_t”,i,sep=””)
data<- data[data$index %in% frequencies$index,]
data$Timestep<- i
data$Year<- unique(MyData$trend$Time)[i]
data$index2<- paste0(data$Species, data$Year)
assign(paste(“spec_pres_t”,i,”real”,sep=””),data)
}

## housekeeping: remove the obsolete full-sized dataframes
rm(spec_pres_t1,spec_pres_t2,spec_pres_t3)

### 3. Calculate Probabilities according to Bijlsma, 2013

for (i in 1:n_timesteps){
data<- get(paste(“spec_pres_t”,i,”real”,sep=””))
trendfacts<- get(paste(“trendfacts_”,i,sep=””))

data$prob<- 1-exp(-(-log(1-frequencies$Freq1[match(data$index,frequencies$index)])* trendfacts$TFactor[match(data$index2,trendfacts$index)]))

assign(paste(“spec_pres_t”,i,”real”,sep=””),data)
}

### 3.1. Make a single dataframe for all timesteps (here: 3)

spec_pres_time<- rbind(spec_pres_t1_real,spec_pres_t2_real,spec_pres_t3_real)

## housekeeping: remove obsolete data frames
rm(spec_pres_t1_real,spec_pres_t2_real,spec_pres_t3_real)

saveRDS(spec_pres_time,”#YourDataOutputFolder#/spec_pres_time.rds”)

### 4. Make the actual graphs
### 4.1. Download shapefile of the country of interest (or use the respective outlines
## from e.g. the world2Hires dataset from the maptools package
```

```
## Here, as an example we present the took the German Bundesländer predefined in an external rds file

bundeslander <- readRDS("./bundeslander.rds")

## Make sure that the CRS of the shapefile is identical to the coordinate CRS in the dataframe ## of species presences

##For any given Species i
  specname<- i
  df<- spec_pres_time[spec_pres_time$Species==i,]
  plot_OP_species <- ggplot(data=df) + geom_sf(data= bundeslander, fill="seashell1")+
                     geom_point(data=df, aes(x=lon, y=lat, color= prob), shape=15)+
                   geom_sf(data= bundeslander, fill=NA) +
                   scale_color_viridis(option=”D”) +
                   facet_wrap( ~ Timestep) +
                   ggtitle(label = i) +
                   theme(plot.subtitle = element_text(hjust = 0.5))

#### END of Script
```

2. **Technical details II: Spatio-temporal Analyses using INLA**
   2.1. **Defining the spatial component**

Here, we outline the spatio-temporal analyses carried out for the analyses presented in the main text. We closely orient on the approach by Zuur & Ieno, (2017) using the R-package INLA (Rue & Martino 2009). R code will be indicated by gray background.

2.2. **Defining a mesh**

**NOTE:** For the present script, an external shapefile depicting the outlines of Germany is necessary. This can e.g. be extracted from the world2Hires dataset of the maptools (Bivand & Lewin-Koh 2018) package. In this case UTM coordinates are necessary for the computation. So, make sure the shapefile is in WGS84, UTM32. INLA cannot cope with spatial files in the simple features format provided by the package ‘sf’ (Pebesma 2018). Make sure to use rgdal (Bivand et al. 2018) for the shapefile here!

library(rgdal)

library(INLA)

\## 1. Read in data

AnalysisDat<- readRDS("./#YourFolderWithData#/#YourDataSetWithSpeciesPresenceEstimates#.rds")

Germany<- readOGR("./Deutschland.shp") # or any other source for a shapefile of Germany

The triangulated spatial mesh that will be used to construct the spatial proximity correlation matrix for u_j_ (see main text) is constructed using an empirical rule of thumb presented by Bakka (2018, <https://haakonbakka.bitbucket.io>). It suggests to specify the maximum edge length of vertices in the triangulated mesh to be defined as 1/5^th^ of the distance of approx. 15% of the observations in the data. The latter can also be specified as the expected range of the spatial autocorrelation (i.e. the expected distance where the spatial autocorrelation is diminishing, Zuur & Ieno 2017).

Figure A5 shows the mesh used in the analyses of Eichenberg et al. (2020). The mesh has 5951 vertices and allows for an extension of a coarser resolution outside of the country boundaries of Germany to account for possible edge distortions.

\## Extract locations from Data

locations<- cbind(AnalysisDat$X_UTM, AnalysisDat$Y_UTM)

exp.range<- round(as.numeric(quantile(dist(locations, upper=T, diag=F)/1000,0.15)),0)

MaxEdge <- exp.range/5

\### Define mesh

mesh <- inla.mesh.2d(boundary = Germany,loc=locations,

max.edge=c(1,5)*(MaxEdge*1000),

cutoff= (MaxEdge*1000)/5)

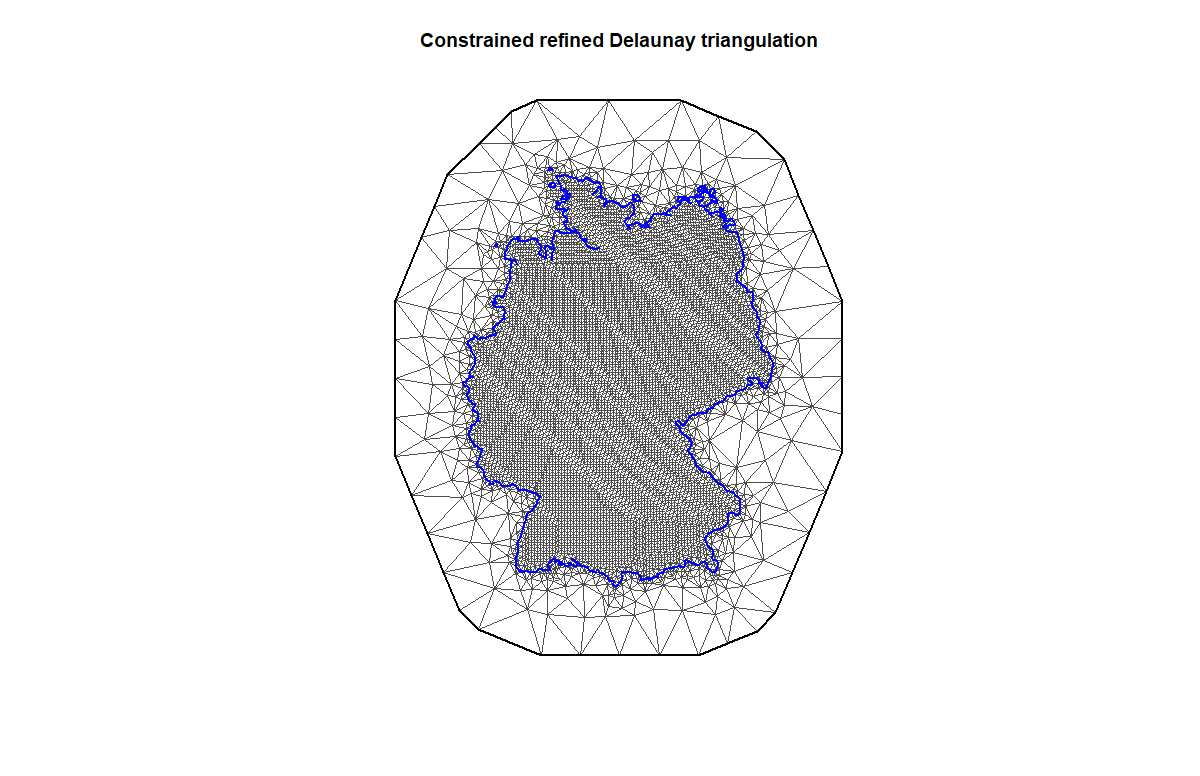

Figure A5: Constrained Delaunay triangulated mesh used in Eichenberg et al (2020). The mesh has 5951 vertices and 11859 triangles. Observations given in the dataset will be placed on or in between the vertices. For further mathematical details see e.g. Zuur & Ieno (2017). The pattern lines seen in the mesh are due to the chosen projection (UTM32).

2.3. **Linking the mesh to the observations in space and time**

To link the observations in the data to the vertices in the mesh, we need to index the data accordingly and tell INLA which index relates to which observation. This is done using a so-called projector matrix. As we need to allow for temporal variation in the spatial field, we also need to indicate, which observation relates to which timestep. This can be done with the “group” argument in the code below:

\## Define index of observations, including the number of study periods (here 3):

n_timesteps<- 3

obs_index <- inla.spde.make.index(name= 'obs', n.spde= spde_sep$n.spde,
 n.group= n_timesteps)

\## Make projector matrix P allowing for temporal variation in the spatial autocorrelation

P <- inla.spde.make.A(mesh,

loc = locations,

group=AnalysisDat$Timestep)

2.4. **Specifying the GMRF**
     2.4.1. **Prior for the range in the spatial autocorrelation**

The specifications for the prior on estimating the spatial autocorrelation in the Model is also based on the empirical rule of thumb by Bakka (2018). The prior itself is specified as a penalized complexity prior (Fuglstad et al., 2019) and is used in the stochastic partial differential equation approach (spde, Lindgren et al., 2011) as the expected distribution of the range in the spatial autocorrelation.

Next, we need to have the observations as well as the predictors needed for the model in a clearly structured data frame.

\### Define the prior for the SPDE

spde <- inla.spde2.pcmatern(mesh,

prior.range=c(exp.range*1000, 0.05), ## expected range in meters

prior.sigma=c(0.5,0.5))	## using a vague prior for the sigma

The objects specified in the code above contain all necessary information to run the spatiotemporal model presented the main text of Eichenberg et al. (2020).

INLA_data<- data.frame(Intercept= rep(1,nrow(AnalysisDat)),

Spec_ric<- AnalysisDat$Spec_ric,

timestep= as.factor(AnalysisDat$Timestep),

grid_cell= as.factor(AnalysisDat$MTB_Q))

2.4.2. **Compile all information to a data stack**

INLA needs all information in a defined format, the data stack. The stack can be created as described below.

\## Compile the data stack

Data_stack <- inla.stack(

tag = "Fit ", ## feel free to name it as you want

data = list(y= INLA_data$Spec_ric), ## the response variable

A = list(1, P), ## the effects in the model,
 ##i.e. predictors below and observations projected to the mesh

effects = list(INLA_data[, c(1,3,4)], ## the predictor variables
 w= obs_index)) ## the index of the observations x mesh

2.4.3. **Specifying the model (Model 3 in main text, Eichenberg et al 2020)**

Finally, the model for the Operation in INLA can be specified as a formula as follows. The argument “rw1” refers to a random walk of order 1, “ar1” refers to a 1^st^ order autoregressive component. They are specified for the temporal component f(timestep) and spatial component f(w, model=spde), respectively. The latter allows for the temporal variation in the spatial component of the model. Finally, the model can be run as follows, including a gamma distribution for the model residuals.

\## Define model formula

f_ar1<- y ~ -1 + Intercept + f(timestep, model=”rw1”) + f(w, model = spde, group=w.group, control.group = list(model="ar1"))

\## Run the model

INLA_ model<- inla(f_ar1,

data=inla.stack.data(Data_stack),

control.predictor = list(

A = inla.stack.A(Data_stack),compute=T, link=1),

control.compute=list(dic=T, waic=T),

family="gamma")

\#### END of Script

NOTE: in Eichenberg et al. (2020, main text) four different subsets of the complete data were fed to the model described above: one comprising the full dataset (all species) and one subset for each species assemblages from the floristic statuses (i.e. natives, archaeophytes, neophytes).

2.5. **Model evaluation of the spatio-temporal models**
     2.5.1. **Spatial fields**

Among-timestep correlations of spatial fields were ρ=0.99 for all models. Strongest bias corrections were applied to archaeophytes and neophytes. Bias correction was also most prominent at the edges of the federal republic as well as in the alpine region and the islands of the northern sea. Moreover, in archaeophytes and neophytes, strong bias correction was necessary in the center of Germany (Figure A6).

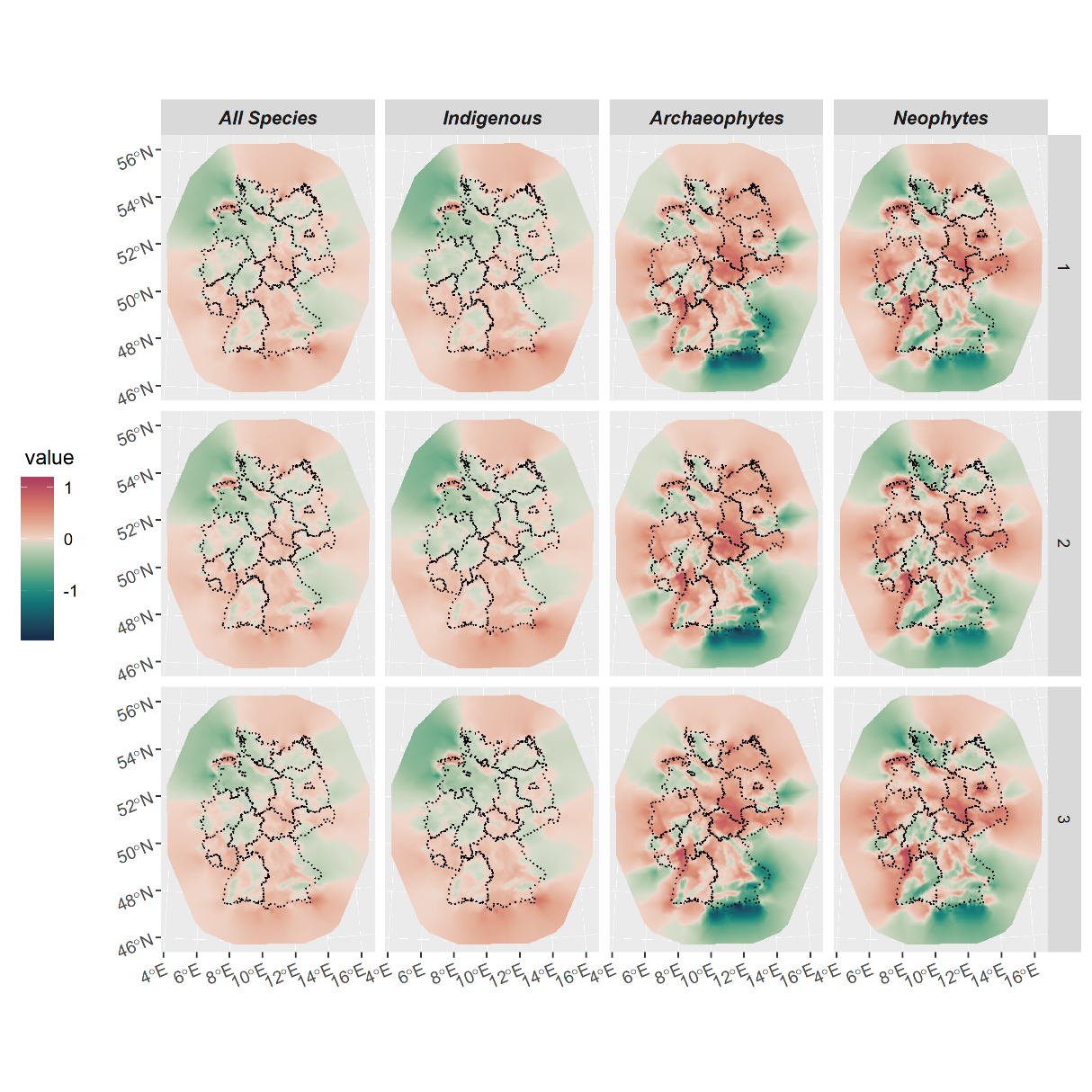

Figure A6: Spatial fields of the spatio-temporal models. Value refers to the value of bias-correction applied to the respective grid to remove spatial autocorrelation. “1”: fist timestep (1960 – 1987); “2”: second timestep (1988 – 1996); “3”: third timestep (1997 – 2017).

Figure A7 shows the distribution of residuals of the spatio-temporal models with and without accounting for the spatio-temporal dependency in the data. The model without accounting for spatio-temporal dependencies shows strong spatial biases in residuals, whereas the model accounting for spatio-temporal dependencies shows much less bias in the residuals. DIC values also indicates a much better fit for the model accounting for these dependencies (lower is better).

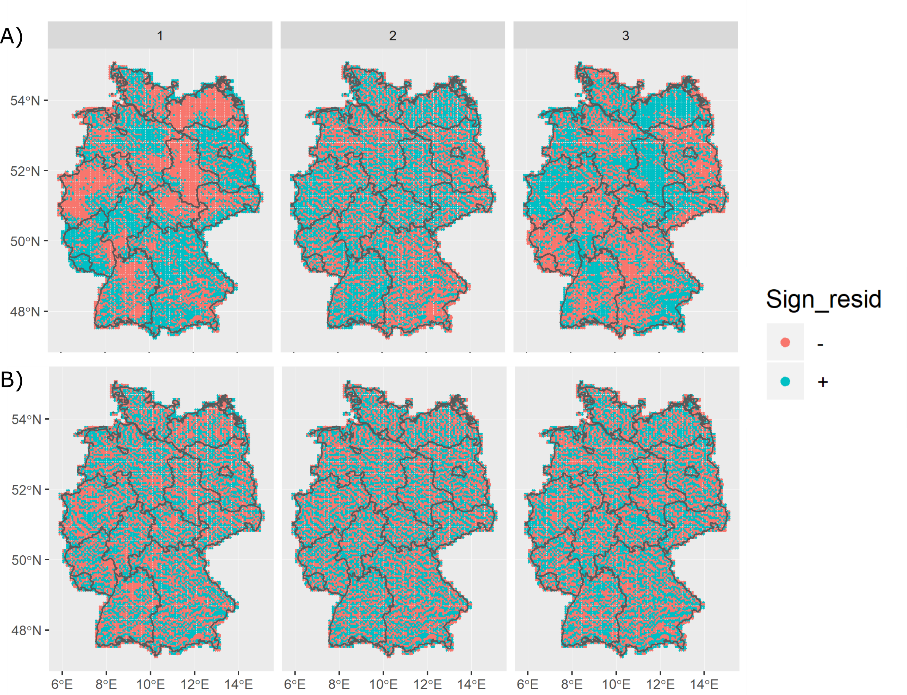

Figure A7: Spatial distribution of residuals in a spatially explicit model of archaeophyte grid-cell species richness without (A) and with (B) temporal variation of the spatial component. Colors indicate the sign of model residuals with salmon= negative and cyan= positive residuals. The model in B performed significantly better than A; DIC criterion (lower is better): DIC (A): 150395.8; DIC (B): 147081.9; other floristic status group models behaved similar.

The mean range of spatial autocorrelation was 155 km, 136 km, 129 km and 169 km for all species, natives, archaeophytes and neophytes, respectively (see Figure A8).
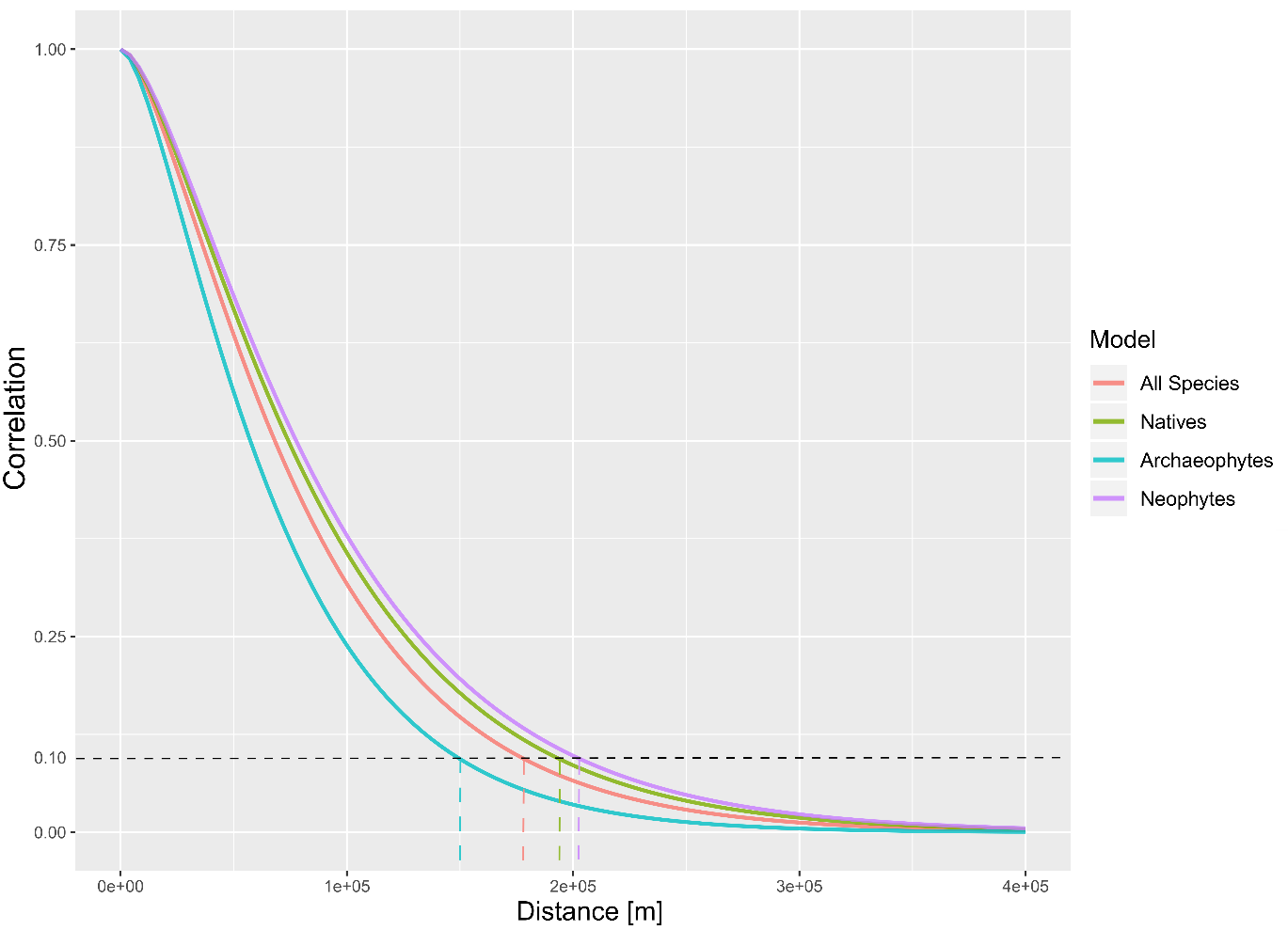

Figure A8: Rangeplot for all models. The range indicates the distance (in meters) from where the spatial autocorrelation diminishes (i.e. Correlation < 0.1, dashed horizontal line). Colors represent values for the different models, according to floristic status assemblages or across all species, respectively. Dashed vertical lines: range estimates. The code to produce this plot is available in the R-package ‘ggregplot’ (gfalbery, <https://github.com/gfalbery/ggregplot/>).

2.5.2. **Residuals in time and space**

Figure A9 shows the distribution of residuals of the spatio-temporal models. These show only very minor patterns of spatio-temporal autocorrelation for the models from all species as well as indigenous species.

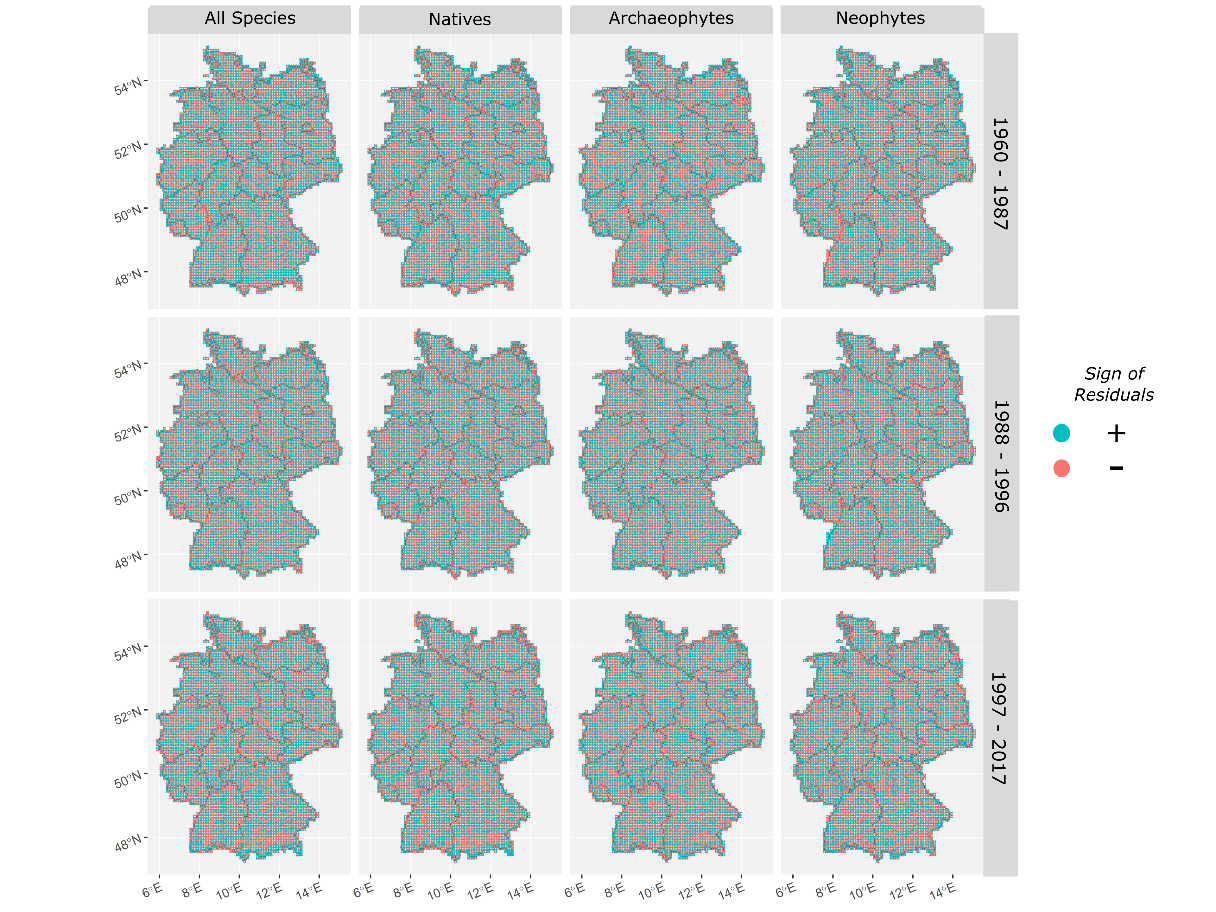

Figure A9: Spatio-temporal distribution of the residuals of the spatio-temporal mixed effects models described above. The optimal case is to see no clustering of residuals in certain regions as well as an even distribution of absolute values of residuals.

Figure A10 shows variograms to assess spatial autocorrelation in the residuals of a model. No signs of residual autocorrelations are found in any of the models.

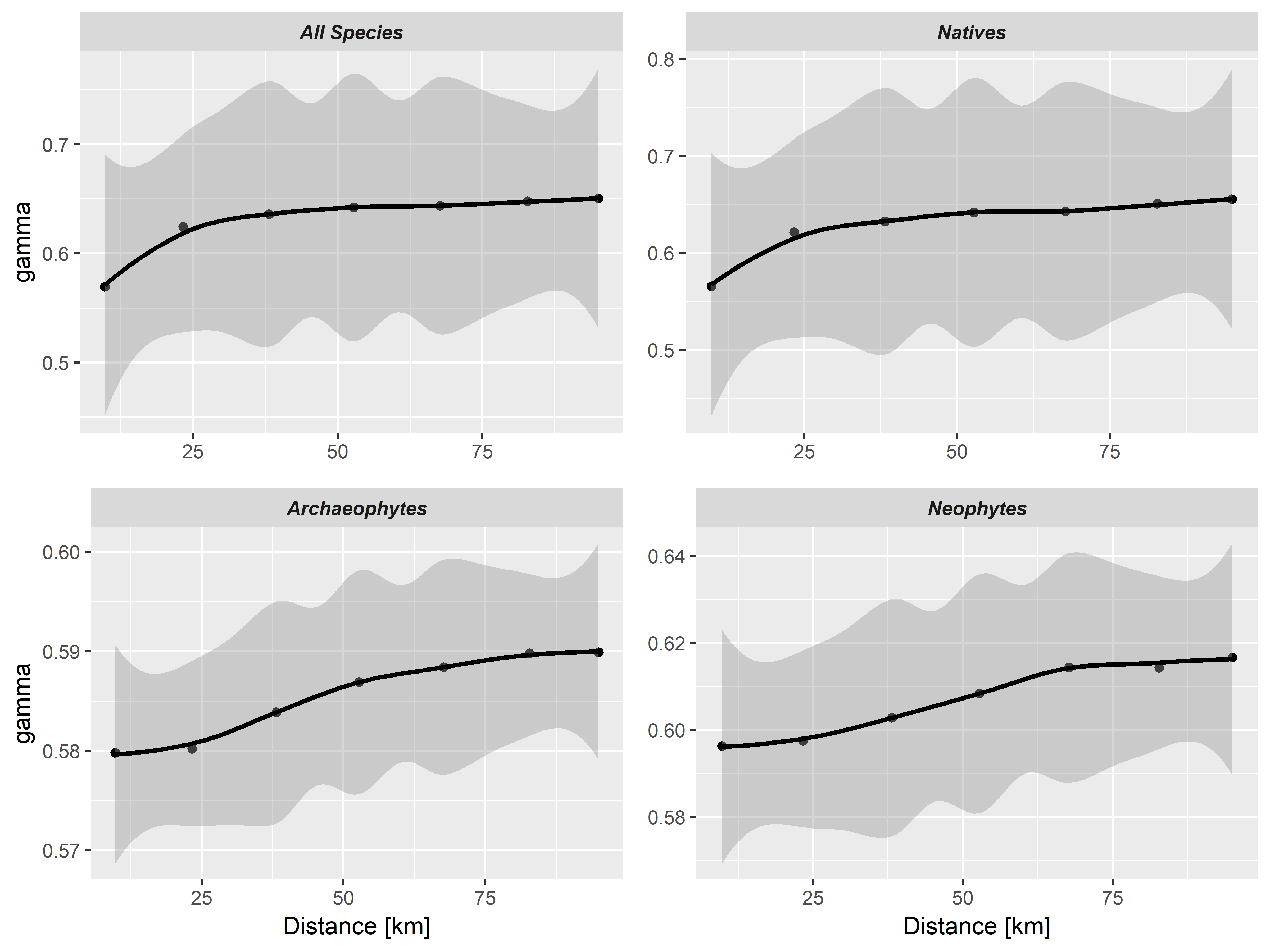

Figure A10: Variograms of the residuals of the spatio-temporal mixed effects models for all floristic statuses, separately. “gamma” is the strength of spatial autocorrelation. Shaded areas are confidence intervals according to a loess smoother. A straight line indicates no spatial autocorrelation in the residuals of a model; uncertainty estimates point towards no significant residual spatial correlation in the models.
